# Distinct interhemispheric connectivity at the level of the olfactory bulb emerges during *Xenopus laevis* metamorphosis

**DOI:** 10.1101/2021.05.11.443461

**Authors:** Lukas Weiss, Paola Segoviano Arias, Thomas Offner, Sara Joy Hawkins, Thomas Hassenklöver, Ivan Manzini

## Abstract

The olfactory system of anuran tadpoles requires substantial restructuring to adapt to the lifestyle of the adult frogs. *Xenopus laevis* tadpoles have a single main olfactory epithelium in the principal nasal cavity associated with aquatic olfaction. After metamorphosis, this epithelial surface is transformed into the adult air-nose and a new epithelium, the adult water-nose, is present in the middle cavity. Impacts of this massive remodeling on odor processing, behavior and network structure are still unexplored.

In the present study, we used neuronal tracings, calcium imaging and a behavioral assay to examine the functional connectivity between the epithelium and the main olfactory bulb during metamorphosis. In tadpoles, olfactory receptor neurons in the principal cavity epithelium project axons to glomeruli in the ventral main olfactory bulb. During metamorphosis, these projections are gradually replaced by receptor neuron axons emerging from the newly forming middle cavity epithelium. Despite this metamorphotic reorganization in the ventral bulb, two spatially and functionally segregated odor processing streams remain undisrupted. In line with this, metamorphotic rewiring does not alter behavioral responses to waterborne odorants. Contemporaneously, newly formed receptor neurons in the remodeling principal cavity epithelium project their axons to the dorsal part of the bulb. The emerging neuronal networks of the dorsal and ventral main olfactory bulb show substantial differences. Glomeruli around the midline of the dorsal bulb are innervated from the left and right nasal epithelia. In addition, postsynaptic projection neurons in the dorsal bulb predominantly have smaller tufts and connect to multiple glomeruli, while more than half of projection neurons in the ventral bulb have a single, bigger tuft.

Our results show that during the metamorphotic reconstruction of the olfactory network the ‘water system’ remains functional. Differences of the neuronal network of the dorsal and ventral olfactory bulb imply that a higher degree of odor integration takes place in the dorsal main olfactory bulb. This is likely connected with the processing of different odorants, airborne vs. waterborne, in these two parts of the olfactory bulb.

## Introduction

Tadpoles of most anuran amphibians share a similarly structured olfactory periphery, consisting of the main olfactory epithelium in the principal nasal cavity (PC), a vomeronasal organ (VNO), as well as some minor additional epithelial surfaces (Jungblut et al., 2021; Weiss et al., 2021). This is well documented for all major groups of anurans: Archaeobatrachians (Benzekri and Reiss, 2012), Mesobatrachians (Manzini and Schild, 2010) and Neobatrachians (Jermakowicz et al., 2004; Jungblut et al., 2011, 2017; Nowack and Vences, 2016; Quinzio and Reiss, 2018).

The transition from an aquatic tadpole to a more or less water-independent adult frog requires the neuronal network associated with odor processing to restructure and rapidly adapt to the new habitat (Duellman and Trueb, 1994; Wells, 2007; Reiss and Eisthen, 2008). During metamorphosis, the olfactory periphery transforms into a tripartite chamber system, consisting of the main olfactory epithelium in the PC, a middle cavity (MC) lined with non-sensory epithelium in most species, and the VNO (Helling, 1938; Reiss and Eisthen, 2008). While the larval system (PC and VNO) is specialized for the detection of waterborne odors, the sensory epithelium in the adult PC is associated with sensing volatile odors (for reviews, see Reiss and Eisthen, 2008; Weiss et al., 2021).

In contrast to many other anurans, adults of the pipid frog *Xenopus laevis* are fully water-dwelling (Wells, 2007; Reiss and Eisthen, 2008) and only occasionally move overland (Du Plessis, 1966; Measey, 2016). The special ecology of adult *Xenopus* is reflected in the presence of a specialized ‘water-nose’ in the MC, which starts to form around the premetamorphotic stage 51 after Nieuwkoop and Faber (Föske, 1934; Nieuwkoop and Faber, 1994; Reiss and Burd, 1997b, 1997a; Hansen et al., 1998; Higgs and Burd, 2001; Dittrich et al., 2016). The epithelium in the adult MC exhibits strong similarities with the larval PC. It contains both major types of olfactory receptor neurons (ORNs, ciliated and microvillous; Hansen et al., 1998), is responsive to common waterborne olfactory stimulants like amino acids (Sorensen and Caprio, 1998; Syed et al., 2017), and expresses a similar set of olfactory receptors (Freitag et al., 1995, 1998; Amano and Gascuel, 2012; Syed et al., 2013, 2017).

During metamorphosis, major remodeling occurs in the PC of larval *Xenopus* caused by massive cell death and replacement of ORNs (Hansen et al., 1998; Higgs and Burd, 2001; Dittrich et al., 2016). The remodeled adult PC is eventually composed of only ciliated ORNs and expresses olfactory receptors putatively responsive to airborne odorants (Freitag et al., 1995; Mezler et al., 1999, 2001), thus assuming the role of the adult ‘air nose’ (Föske, 1934; Hansen et al., 1998; Syed et al., 2017). In contrast, the VNO does not seem to change significantly during metamorphosis in regard to its cellular composition or function (Hansen et al., 1998; Dittrich et al., 2016).

The functional and cellular segregation of the olfactory periphery also translates to the level of the olfactory bulb. Receptor neurons in the VNO project their axons towards their target structures, the glomeruli, in the accessory olfactory bulb (Reiss and Eisthen, 2008; Jungblut et al., 2012). ORNs residing in the PC of larval *Xenopus* project their axons to glomeruli arranged in the ventral portion of the main olfactory bulb (vMOB) (Reiss and Burd, 1997a; Weiss et al., 2020a). During metamorphosis, these axonal projections from the PC to the vMOB are replaced by ORN axons originating in the newly formed MC, the adult ‘water nose’ (Key and Giorgi, 1986; Hofmann and Meyer, 1991; Franceschini et al., 1992; Reiss and Burd, 1997a). Newly generated ORNs in the PC are projecting their axons towards the dorsal part of the main olfactory bulb (dMOB) during metamorphosis (Hofmann and Meyer, 1991; Reiss and Burd, 1997a; Gaudin and Gascuel, 2005). Since it is solely innervated by ORNs residing in the adult ‘air-nose’, the dMOB is putatively associated with the processing of volatile odorants (Föske, 1934; Reiss and Eisthen, 2008; Weiss et al., 2021).

Functionally, the axonal projections to the vMOB of *Xenopus* tadpoles have been shown to be segregated into two major odor processing streams: projections to laterally located glomeruli use a second messenger cascade dependent on phospholipase C and diacylglycerol (DAG), while projections to medially located glomeruli are dependent on the adenylate cyclase and use cyclic adenosine monophosphate (cAMP) as a second messenger (Gliem et al., 2013; Sansone et al., 2014). A substantial portion of the lateral glomeruli are responsive to amino acids and putatively innervated by microvillous ORNs (Gliem et al., 2013). Medially located glomeruli are putatively connected to ciliated ORNs and mainly respond to alcohols, aldehydes and amines instead (Gliem et al., 2013). While there is evidence, that the anatomy of the glomeruli in the vMOB remains stable during metamorphosis (Gaudin and Gascuel, 2005; Weiss et al., 2020a), it is still unknown whether the innervation shift from the larval PC to the adult MC affects its function and/or its behavioral output.

Odor information transferred to the glomeruli in both parts of the MOB is subsequently relayed to the postsynaptic cells, the projection neurons (Imamura et al., 2020). In amphibians, these neurons feature one or more apical dendrites that terminate in densely branched tufts within glomeruli, and an axon projecting to higher brain centers (Jiang and Holley, 1992a; Dryer and Graziadei, 1994; Nezlin et al., 2003; Imamura et al., 2020). It has been shown in other vertebrates that the morphology of projection neurons varies based on their function or their association with distinct olfactory subsystems (Nagayama et al., 2014; Imamura et al., 2020; Braubach and Croll, 2021). In amphibians, such analyses are lacking so far.

In the present work, we analyze the neuronal network restructuring of the MOB during metamorphosis on an anatomical, functional, and behavioral level. We show that ORN axonal projections from the larval PC to the vMOB are progressively replaced by ORN axons originating in the newly formed MC. Despite these massive reorganization processes, the functional segregation into two separate odorant processing streams in the vMOB remains stable. Similarly, the remodeling does not change behavioral responses to amino acids, implying that connections to higher brain centers remain intact during the process. Axons from newly developing ORNs in the remodeled PC project to the dMOB, forming an unpaired structure around the interhemispheric midline. We show that the dMOB network follows a wiring logic distinct from the vMOB, with a progressively higher number of ORN axons projecting to glomeruli in the contralateral hemisphere. Additionally, the population of postsynaptic projection neurons of the vMOB and the dMOB differ morphologically, revealing a higher prevalence of multi-tufted neurons connected to multiple glomeruli associated with the dMOB. These features point towards a higher degree of integration between the hemispheres and across several glomeruli in the dMOB, possibly adaptive to the processing of volatile odorants. The processing of waterborne odorants in the postmetamorphotic vMOB on the other hand seems to mirror the larval system.

## Material and Methods

### Animals and tissue preparation

All animals used in this study were wild type or albino *Xenopus laevis* (both sexes), kept and bred at the University of Giessen at a water temperature of 20°C in water tanks with constant water circulation. Developmental stages were defined according to Nieuwkoop and Faber (Nieuwkoop and Faber, 1994). Before experimental procedures, the animals were anesthetized using 0.02% MS-222 (ethyl 3-aminobenzoate methanesulfonate; TCI Germany) in tap water. For tissue preparation, anesthetized animals were killed by severing the transition between the brainstem and spinal cord. For lower staged animals, a tissue block containing the noses and the rostral part of the telencephalon was removed. For higher staged animals, the olfactory nerve was cut close to the noses and the entire brain was taken out of the cartilage. All animal procedures were performed in accordance with the guidelines of Laboratory animal research of the Institutional Care and Use Committee of the University of Giessen (V 54 – 19 c 20 15 h 01 GI 15/7 Nr. G 2/2019; 649_M; V 54 – 19 c 20 15 h 02 GI 15/7 kTV 7/2018).

### Tracings of olfactory receptor neurons via fluorophore-coupled wheat germ agglutinin (WGA) and electroporation

For bulk loadings of olfactory projections from the nose to the MOB, we filled the olfactory epithelia with droplets of approx. 3 μl of fluorophore-coupled wheat germ agglutinin (WGA Alexa Fluor 488 or 594 conjugate, Thermo Fisher) diluted at a concentration of 10 mg/ml in saline Frog Ringer (in mM: 98 NaCl, 2 KCl, 1 CaCl_2_, 2 MgCl_2_, 5 glucose, 5 Na-pyruvate, 10 Hepes, pH 7.8). For bulk electroporations, we placed dried dye crystals of fluorophore-coupled dextrans (Alexa Fluor dextran 488, 594, Cascade Blue dextran, Thermo Fisher or Cal520 dextran conjugate, AAT Bioquest, 10kDa, 3 mM in Frog Ringer) in the nostrils and applied six electric square pulses using two platinum electrodes (15 V, 25 ms duration at 2 Hz with alternating polarity) to each nostril (for detailed protocol see Weiss et al., 2018). After the procedures, animals were left to recover for at least 24 hours before tissue preparation and imaging.

### Sparse cell electroporation in the olfactory epithelium and the olfactory bulb

We sparsely electroporated ORNs in the nasal epithelia of the PC/MC and projection neurons in the vMOB/dMOB using micropipettes pulled from borosilicate glass capillaries (Warner instruments, resistance 10-15 MΩ) filled with fluorophore-coupled dextrans (Alexa Fluor dextran 488 and 594; 3 mM in Frog Ringer). The dye-filled micropipettes were mounted on the headstage of a single cell electroporator (Axoporator 800A; Axon Instruments, Molecular Devices) equipped with a wire electrode and approached cells in the olfactory epithelium (PC/MC) or the MOB using a micromanipulator. A 500 ms train of square voltage pulses (50 V, single pulse duration 300 μs at 200– 300 Hz) was applied.

For ORN labeling, the animals were first anesthetized and sparse cell electroporation was repeated at multiple locations using Alexa Fluor 488 dextran in the MC and Alexa Fluor 594 dextran in the PC to trace their respective projections during metamorphosis. After the procedure, animals were left to recover for at least 24 hours prior to image acquisition (Hassenklöver and Manzini, 2014). Sparse labeling of projection neurons in the MOB was conducted in an excised tissue block containing the olfactory system. We fixed the bulb and the caudal portion of the olfactory nerve under a platinum grid stringed with nylon threads, approached the micropipette to the cell layer containing MOB projection neurons, and applied the voltage pulse trains as described above (detailed protocol Weiss et al., 2018).

### Image acquisition and processing of morphological images

Images were acquired as multi-color virtual image stacks with a z-resolution of 1–3 μm using multiphoton microscopy (Nikon A1R-MP) at an excitation wavelength of 780 nm. We used ImageJ (Schindelin et al., 2012; RRID:SCR_003070) to adjust the brightness and contrast of the image stacks and applied a median filter to remove pigmentation-derived artifacts in some images. Separate images were stitched together where necessary (Preibisch et al., 2009). For thresholding analyses conducted on the image-stacks, we eliminated tissue-derived autofluorescence by subtracting the blue-wavelength color channel (when no blue-emitting dye was introduced into the tissue). Images are presented as maximum intensity projections along the z-axis or in 3D using the 3D-viewer implemented in ImageJ.

### Functional calcium imaging and data processing

For functional calcium imaging in the vMOB, we loaded the ORNs with a calcium sensitive dextran coupled dye (Cal 520 dextran conjugate, 10 kDa, AAT Bioquest; 3 mM in Frog Ringer) via bulk electroporation (Weiss et al., 2018) as described above. After killing the animal, we cut out the tissue block containing the noses and the anterior part of the brain and removed the ventral palatial tissue as well as tissue around the sensory epithelia to facilitate odorant flow into the nasal cavities. The tissue block was positioned on the stage of the multiphoton microscope using a platinum grid stringed with nylon threads. A perfusion manifold with silicone tubing outlet (Milli Manifold; ALA Scientific) connected to a gravity-fed multi-channel perfusion system (ALA-VM-8 Series; ALA Scientific) was positioned in front of the nasal cavity and a constant Ringer flow was established. Ringer was constantly removed from the recording chamber via a syringe needle connected to a peristaltic pump via silicone tubing. Fast volumetric recordings of the ORN axon terminals in the glomeruli of the vMOB were made using the resonant scanning mode of a multiphoton microscope (780 nm excitation wavelength). We measured time series of 3D virtual image stacks (lateral dimensions: 509 × 509 μm, 512 × 512 pixel ; axial dimensions: 180–300 μm, inter-plane distance 4–7 μm) at 0.5–1 Hz per image stack (Offner et al., 2020). Stimuli were applied with a duration of 5 seconds and an inter-stimulus interval of 60 seconds and repeated at least twice. The stimuli used were: Amino acid mixture: L-valine, L-leucine, L-isoleucine, L-methionine, glycine, L-serine, L-threonine, L-cysteine, L-arginine, L-lysine, L-histidine, L-tryptophan, L-phenylalanine, L-alanine, L-proline (100 μM in Frog Ringer solution); Amine mixture: 2-phenylethylamine, tyramine, butylamine, cyclohexylamine, hexylamine, 3-methylbutylamine, N,N-dimethylethylamine, 2-methylbutylamine, 1-formylpiperidine, 2-methylpiperidine, N-ethylcyclohexylamine, 1-ethylpiperidine, piperidine (100 μM in Frog Ringer solution); Bile acid mixture: taurocholic acid, cholic acid, glycholic acid, deoxycholic acid, (100 μM in Frog Ringer solution); Odorant mixture: mixture of amino acids (excluding proline and alanine), amines, bile acids, alcohols and aldehydes (positive control, 50 μM in Frog Ringer solution) (Gliem et al., 2013); Frog Ringer solution (negative control); Forskolin, a direct stimulant of the adenylate-cyclase and the cAMP second messenger pathway. Forskolin was dissolved in DMSO (stock of 10 mM) and used at a final concentration of 100 μM. All chemicals were purchased from Sigma Aldrich. The mixtures were stored as frozen aliquots and diluted to their final concentration shortly before the experiments.

Data evaluation was done using Python. A piecewise-rigid motion correction algorithm was applied to remove motion artifacts (Pnevmatikakis and Giovannucci, 2017). We used the CaImAn toolkit for calcium imaging data to generate denoised and deconvolved templates of the 3D image stacks (Pnevmatikakis et al., 2014, 2016; Friedrich et al., 2017; Giovannucci et al., 2019). These templates were used for maximum intensity projections along the z-axis of fluorescence intensity difference maps. Intensity difference maps resulted from the difference between the peak fluorescence intensity of post-stimulus responses (averaged from three timeframes) and baseline fluorescence prior to stimulus onset (averaged from five frames). Individual timeframes had a duration between 1.5–2 seconds. Responding regions were selected according to the following criteria: i) Regions had to be bigger than 100 pixels in area. ii) They had to be reactive to at least one applied stimulus with a stimulus-evoked fluorescence intensity amplitude exceeding the average of all median values calculated for the amplitudes of each responding region in the dataset. iii) The deconvolved timeseries generated by CalmAn had to pass an additional quality control, comparing them to the raw traces and discarding regions outside the boundaries of the glomerular clusters (using an interactive custom written Python script).

Changes in fluorescence of responsive regions over time were baseline corrected using asymmetric least squares smoothing (Eilers and Boelens, 2005) and response tuning profiles were determined by normalizing each individual fluorescence intensity timeseries by its maximum intensity value (baseline:0, maximum:1). We then defined a threshold below which to ignore amplitude peaks in the expected peak response intervals (12.5%). The tuning profiles of responding regions were then defined by the set of stimuli that led to response amplitudes above 12.5% of the maximum amplitude of the respective time series.

For the spatial analysis of response profiles, responsive regions were first sorted according to their tuning profiles into forskolin-responsive (fsk+) and non-responsive (fsk-) regions. Both groups of regions were subcategorized based on their further response profiles to the applied odorant mixtures. A region’s position within the glomerular clusters was measured in relation to manually defined points at the medial-lateral and anterior-posterior edges of the ORN axonal projections in each individual vMOB. Along these two axes, we estimated the density of responsive regions using Gaussian kernel density estimates. Scott’s rule implemented in the Python library SciPy (https://scipy.org/) was used to calculate the estimator bandwidth (Scott, 1992; Virtanen et al., 2020).

### Analysis of ORN projections to the MOB

The volumes of the ventral and dorsal portions of the ORN projections in the MOB (presented in Figure 1) were manually annotated based on their visual outlines in the image stacks and measured using the Segmentation Editor implemented in ImageJ (Schindelin et al., 2012). They were subsequently analyzed using Python. The data show the percentual share of ORN projections in the dMOB and the vMOB relative to the volume of the entire ORN projections (dMOB + vMOB). Projections in both hemispheres were summed up for this analysis.

**Figure 1.**
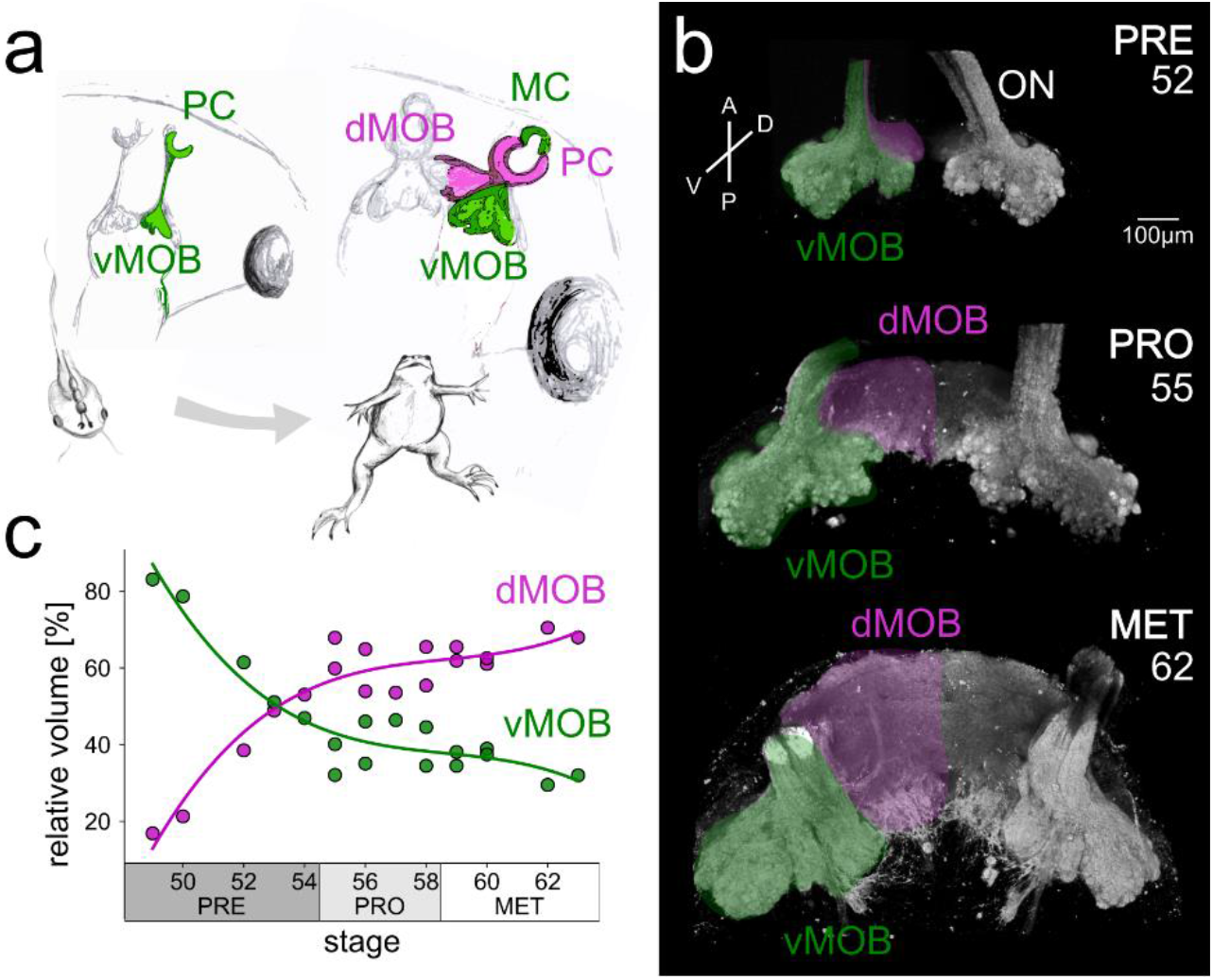
Differential development of ORN projection areas in the ventral and dorsal MOB during metamorphosis. **(a)** The scheme shows axonal projections of ORNs residing in the PC innervating the vMOB (green) in the tadpole. After metamorphosis, the newly formed MC connects to the vMOB (green), while the remodeled PC epithelium innervates the dMOB (magenta). **(b)** 3D projections of the caudal part of the ONs and the ORN axonal projections during premetamorphosis (stages up to 54), prometamorphosis (55–58) and metamorphosis proper (59–65), stained with fluorophore-coupled WGA. Numbers in the panels indicate the developmental stages of the depicted animals. The projections to the vMOB are already present in premetamorphosis and retain their morphology during development (green). The dMOB projections start to form around stage 50 and increase dramatically in size until the end of metamorphosis (magenta). In contrast to the vMOB, the dMOB is not divided at the interhemispheric midline in later staged animals. **(c)** Volume share of axonal projections in the vMOB (green) and dMOB (magenta) relative to the total axonal projections during metamorphotic development. Around stage 50, the dMOB only occupies less than 20% of the total axonal projection volume and grows to occupy more than 60% around stage 62. The relative growth of the dMOB is best described by a cubic curve (R^2^ = 0.9). A anterior, D dorsal, dMOB dorsal main olfactory bulb, MC middle cavity, MET metamorphosis proper, ON olfactory nerve, P posterior, PC principal cavity, PRE premetamorphosis, PRO prometamorphosis, V ventral, vMOB ventral main olfactory bulb.

For the analysis of MC and PC projections to the MOB (shown in Figure 2), we separated image planes containing the vMOB and the dMOB and subsequently reduced the two image stacks to maximum intensity projections along the z-axis. Images were binarized using the Maximum Entropy thresholding method implemented in ImageJ (Kapur et al., 1985). Projections from the MC and PC were detected in two separated color channels. The percentual share of pixels containing ORN fibers from the MC and PC was calculated in relation to the total number of pixels containing fluorescent signal in either channel. This analysis was conducted separately for the dMOB and the vMOB.

**Figure 2.**
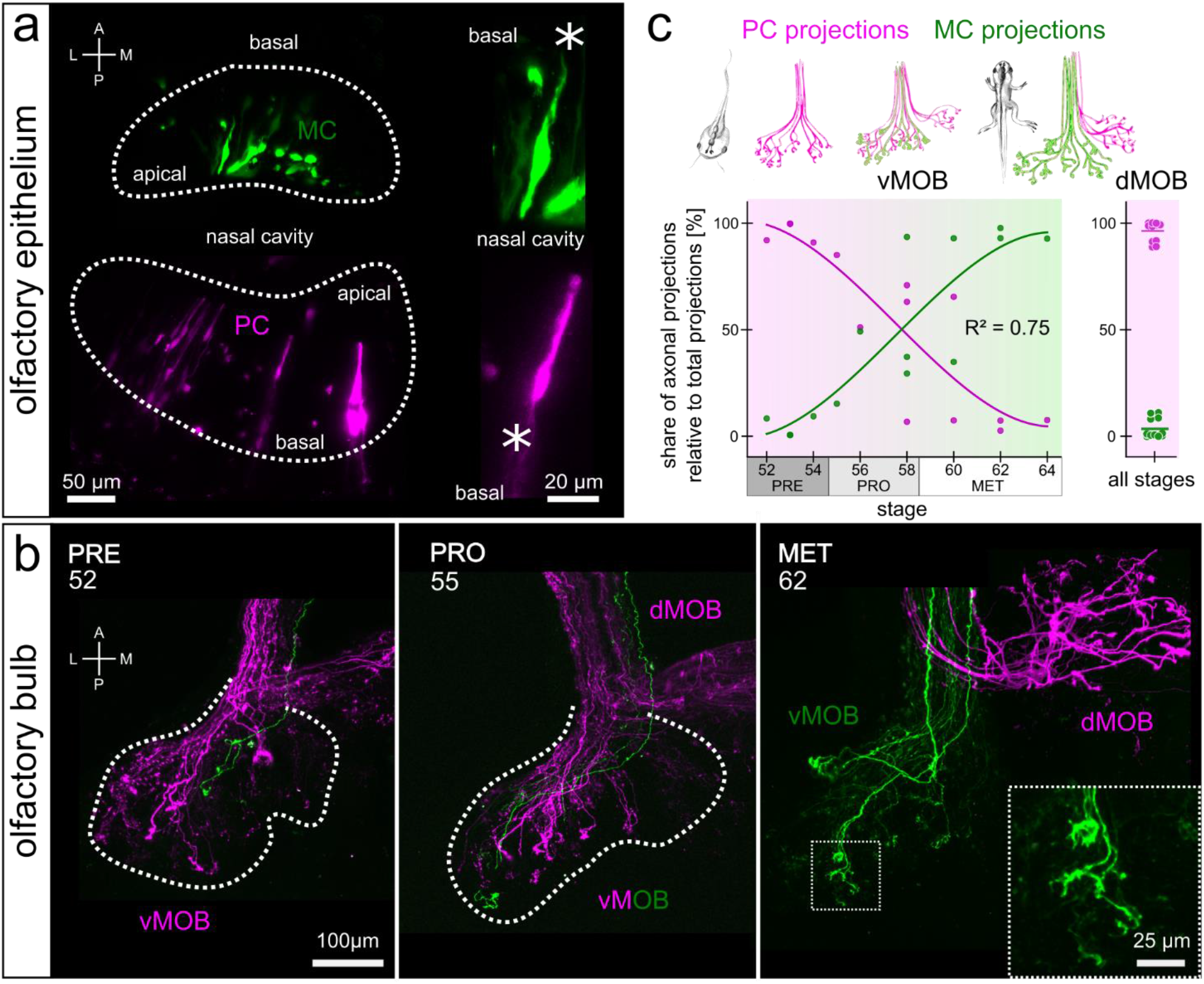
MC axonal projections gradually replace PC fibers in the vMOB during metamorphosis. New PC neurons innervate the dMOB. **(a)** Labeling of ORNs in the olfactory epithelia of MC (green) and PC (magenta) using different fluorophore-coupled dextrans via sparse cell electroporation. Dotted lines indicate the boundaries of the sensory epithelia. Close-ups show single bipolar ORNs extending their dendrite into the nasal cavity. Asterisks highlight the axons. **(b)** Tracings of axonal projections in the vMOB of tadpoles during different developmental stages (indicated by numbers in the panels). During premetamorphosis, the first ORN axons originating in the newly forming MC reach the glomerular layer (green, left). In prometamorphotic animals, both MC (green) and PC (magenta) axons are innervating glomeruli in the vMOB (middle). During metamorphosis proper, the vMOB is solely innervated by axons originating in the MC (green), the newly formed dMOB only from PC fibers (magenta, right). Dotted lines indicate the outline of glomerular projections in the vMOB. The close-up shows axonal terminals of MC fibers in the glomeruli of the vMOB. **(c)** Scheme of innervation shift in the vMOB and newly formed connections to the dMOB (top). Percentual share of MC (green) and PC (magenta) axonal projections in the vMOB during metamorphotic development follows a curve of third degree (R^2^ = 0.75). During premetamorphosis, the majority of vMOB projections originate in the PC, while at the end of metamorphosis, almost all axonal projections to the vMOB originate in the MC. The plot on the right shows the same data for the dMOB, which is only innervated by PC axons throughout development. A anterior, dMOB dorsal main olfactory bulb, L lateral, M medial, MC middle cavity, MET metamorphosis proper, ON olfactory nerve, P posterior, PC principal cavity, PRE premetamorphosis, PRO prometamorphosis, vMOB ventral main olfactory bulb.

To calculate the overlap of ORN axonal projections originating in the left and right olfactory epithelia (Figure 5), we applied a median filter with a size of 10 pixels to the virtual two-color channel image stacks in ImageJ and subsequently binarized the image applying Huang’s fuzzy thresholding method (Huang and Wang, 1995). We counted pixels containing signals in both color channels of the image for each image plane of the virtual image stack and calculated their percentual share in relation to the number of all pixels containing any fluorescence signal. Separate analyses for dMOB and vMOB were conducted.

### Reconstruction and analysis of labeled projection neurons

The morphology of projection neurons in the MOB labeled via electroporation was semi-automatically reconstructed from the image stack using Vaa3D (Peng et al., 2010; RRID:SCR_002609). Branching- and endpoints of the projection neurons were defined and translated into a hierarchical 3D tree-structure with the soma of the neuron as the root-point. Each segment of the neuron-tree has a single parent segment that it connects to. Each projection neuron connects to at least one olfactory glomerulus with a dendritic tuft. The number of tufts was determined using the DBSCAN algorithm (Density-Based Spatial Clustering of Applications with Noise) implemented in the scikit learn machine learning package written for Python (Pedregosa et al., 2011; Weiss et al., 2020b). The algorithm classifies an accumulation of end- and branching points of the neuronal structure as tuft-clusters if there are more than 5 points in spatial proximity (<15 μm). All branching- and endpoints that are sparsely distributed in space are classified as blunt endings and not part of a tuft-cluster. The distances of the dendritic tufts to the root of the tree (soma) were measured along the dendritic branches, and the distance between two tufts as Euclidean distance in 3D space. The volumes of the tufts were estimated based on the volume of a convex hull of all branching- and endpoints belonging to a single tuft-cluster (see Supplementary Video 1).

### Behavioral assay

Tadpoles of different developmental stages (stages 47–59) were placed in a 1-liter water tank, which was separated into two areas by a dividing wall along approx. half of its length (Figure 4). All areas were freely accessible to the tadpoles. Tadpoles were left to swim without stimulus application for a 2 h habituation period, and the average time they spent per visit to the two areas at either side of the dividing wall was recorded. After the habituation period, 5 ml of an amino acid mixture (100 μM at the application entry point at a speed of 2 ml/min) was applied through a gravity feed to one of the two areas, while a control stimulus (5 ml water) was simultaneously applied on the other side. The average time the tadpoles spent in the amino acid and water control area per visit was recorded for 20 minutes after their first visit to either area using EthoVision tracking software. Tadpoles that did not enter either side within 40 minutes after stimulus application were not included in the analysis.

### Statistics

Averaged data are presented as mean ± standard deviation. A least-squares based polynomial regression curve of third-degree was fitted through the vMOB/dMOB volume data. A linear regression using the RANSAC (Random sample consensus) algorithm (scikit learn, RRID:SCR_002577; Pedregosa et al., 2011) was used to analyze the left/right projection overlap in the dMOB. Statistical significance was tested using a Mann-Whitney-U test. Behavioral data are presented as median values of all experimental animals, and significance was assessed using Wilcoxon signed-rank test for paired data.

## Results

### The dMOB expands drastically during early metamorphosis

In premetamorphotic larvae (up to stage 54), the olfactory epithelium in the PC innervates the vMOB (Figure 1a, left). After metamorphosis is completed (stage 66), the PC epithelium connects to a newly formed dorsal target area (dMOB; magenta) and the *de novo* formed MC epithelium innervates the vMOB (green, Figure 1a, right). We have monitored the stage-by-stage changes of ORN axonal projections to glomeruli in the MOB using fluorophore-coupled wheat germ agglutinin (WGA) tracing (Figure 1b) and calculated their projection volumes in relation to the overall projection volume (Figure 1c).

In premetamorphotic tadpoles (stages 49–54), ORN axonal projections on the ventral side of the MOB are clearly discernable (green shaded area in Figure 1b), while only a few axonal projections in the dMOB are visible (magenta, Figure 1b). At this stage in development, the two MOB hemispheres are clearly separated. During prometamorphosis (stages 55–58) and metamorphosis proper (stages 59–65), the vMOB axonal projections only slightly increase in size, while the dMOB shows a more significant volume increase (Figure 1b, middle and bottom). Around stages 53/54, the dMOB fuses at the midline and forms a single dorsal projection field.

To understand the relative growth of the two MOB parts, we assessed the percentual share of vMOB and dMOB projections during metamorphotic reorganization (Figure 1c). During premetamorphosis, the projections in the vMOB occupied 64 ± 16% and in the dMOB 36 ± 16% of the entire axonal projections in the MOB (n = 5 animals). The relative volume of the vMOB progressively decreases to 40 ± 6% during prometamorphosis (n = 7) and 35 ± 4% during metamorphosis proper (n = 6). Inversely, the dMOB grows to occupy 64 ± 4% of the entire MOB projection volume in the late metamorphotic group. The best fit model for our data was a polynomial curve of third degree (least squares method, R^2^ = 0.9; Figure 1c). The magenta curve in Figure 1c shows a steep increase in the relative volume of the dMOB from less than 20% at stage 50 to 50% around the end of premetamorphosis (intersection points of the two regression curves, Figure 1c). Subsequently, the dMOB percentage is rising steadily but with a smaller slope until reaching between 60–70% in the late metamorphotic stages.

### The innervation of the vMOB dynamically shifts from the larval PC to the MC until metamorphotic climax

While the glomerular structures of the dMOB are formed *the novo*, the vMOB undergoes major transformation processes during metamorphosis. We sparsely labeled ORNs in the newly forming MC (green, Figure 2a) and the PC (magenta, Figure 2a) and imaged their respective axonal projections in the MOB (Figure 2b). The ORNs labeled were bipolar neurons with a long dendritic shaft on the apical side of the epithelium and an axon leaving the epithelia on the basal side to join the olfactory nerve (asterisks, Figure 2a). A single axon originating in the MC reaching the vMOB is found in a tadpole representative for premetamorphosis (green, Figure 2b, left). Most ORN axons innervating glomeruli in the vMOB originated in the PC (magenta). In a prometamorphotic animal of stage 55, the majority of axonal projections still originates in the PC, while few MC fibers can be seen (Figure 2b, middle). At stage 62, the innervation shift is completed, with all axonal projections to the vMOB originating in the MC. Accordingly, the newly forming dMOB only contains axons originating from the PC (Figure 2b, right).

We quantified this dynamic innervation shift in the vMOB by measuring the percentual share of MC- and PC-originating fibers during metamorphosis (Figure 2c). During premetamorphosis (n = 4 animals), 95 ± 5% of the total innervation of the vMOB originates in the PC. A major shift occurs in prometamorphotic stages (n = 5). Initially (stage 55/56), approx. 70% of the projections originate in the PC, while this value drops to around 30–40% in animals of stage 58. ORN projections from the MC are amounting to around 60–70% at the end of prometamorphosis. The vMOB in later metamorphotic animals is innervated mostly by MC axons (82 ± 27%, n = 5). Around the time of metamorphotic climax (stage 60/61), the majority of PC fibers in the vMOB of the premetamorphotic stages is replaced by new ORN axons from the MC. Contrastingly, the dMOB is only innervated by PC axons across all developmental stages (97 ± 4%, n = 14 animals; Figure 2c, right plot).

### Two parallel odor processing streams remain stable during rewiring of the vMOB

ORN axonal projections from the PC epithelium to glomeruli in the vMOB are gradually replaced by MC projections during metamorphotic remodeling. So far it has remained unexplored, whether this significant rewiring affects odor processing and whether the odor map remains unchanged. We loaded the ORN axons with a calcium-sensitive dye and imaged changes in fluorescent signal in their axonal projections upon stimulation with different odorant mixtures. We first applied a mixture of all odorant mixtures as a positive control and the Ringer solution without odorants as a negative control, then the three individual odorant mixtures (amino acids, bile acids, amines) and forskolin (fsk), an activator of the adenylate cyclase and in consequence of the cAMP-dependent second messenger pathway.

Figure 3a shows the spatial distribution of regions responding to odorants (blue) and forskolin (yellow) in the vMOB of a prometamorphotic (Figure 3a, stage 57, above) and a metamorphotic (stage 59, below) animal. For an approximate 3D representation of the responding regions, we split the entire vMOB (right images, Figure 3a) into ventral (left images) and dorsal (middle images) halves. In the ventral halves of the vMOB of both animals, laterally located glomeruli did not respond to forskolin and exhibited transient calcium increases upon stimulation with odorants (Figure 3a, blue). The majority of glomeruli in the medial portion of the ventral vMOB responded to forskolin application (Figure 3a, yellow) and some regions additionally responded to the odorant mixtures (blue and yellow overlay). In the dorsal halves of the vMOBs of both animals, forskolin responsive regions are not confined to the medial side, but were distributed across the entire span of the vMOB. This spatial response pattern was consistent across the 11 analyzed vMOB hemispheres from 8 different animals, and unchanged between prometamorphotic (n = 6 vMOBs, 4 different animals) and metamorphotic animals (n = 5 vMOBs, 4 different animals).

**Figure 3:**
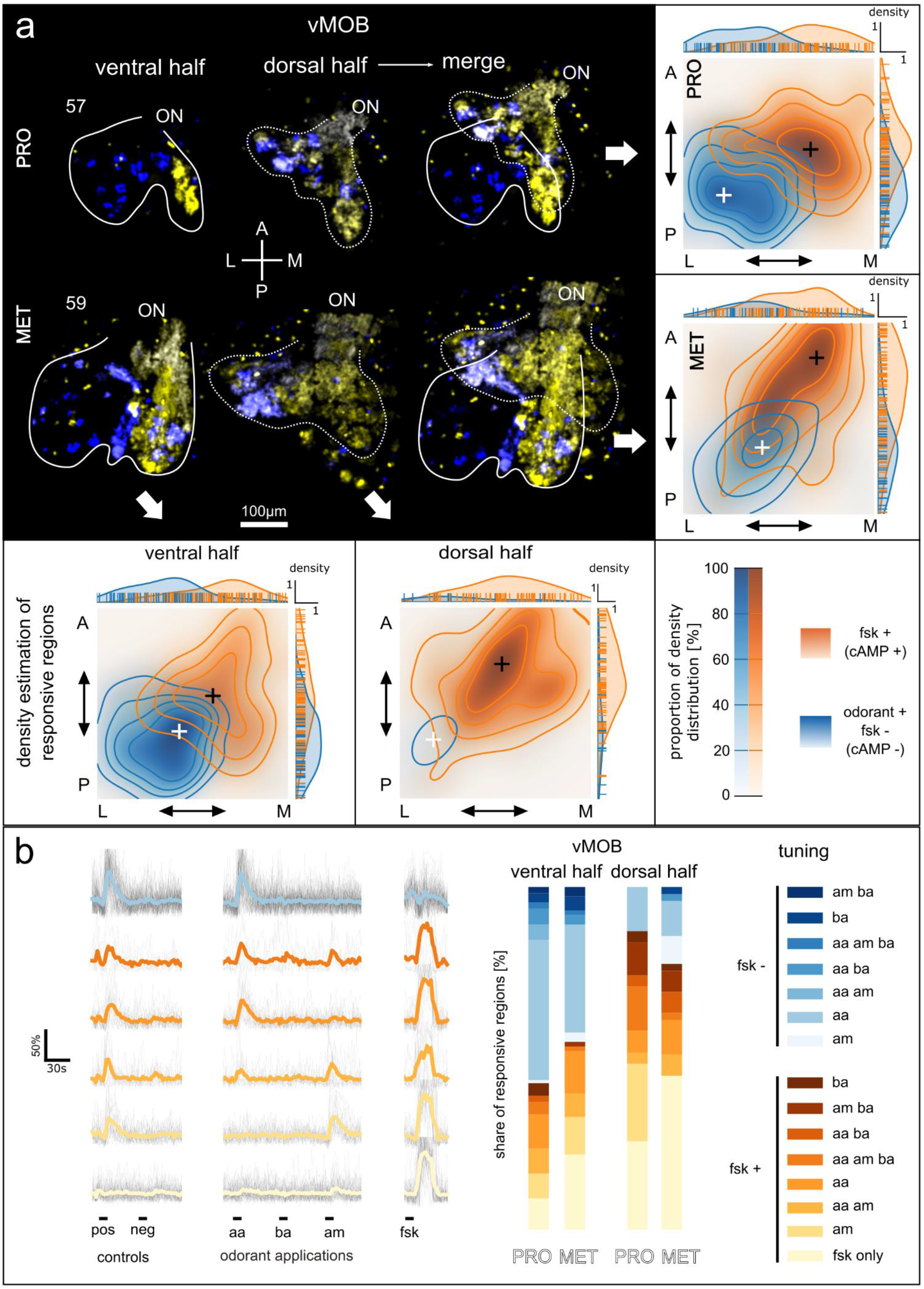
Segregated odorant processing streams in the vMOB remain stable during metamorphotic remodeling. **(a)** Maximum intensity projections of difference maps showing the increase in fluorescence signal after application of the odorant mixture (including amino acids, amines and bile acids; blue) and forskolin (yellow). The ventral (left) and dorsal (middle) halves of the vMOB of a prometamorphotic (stage 57, above) and a metamorphotic animal (stage 59, below) are shown. The two images to the right show the merged ventral and dorsal halves of the depicted vMOBs. White solid and dotted lines indicate the outline of ORN axonal projections of the ventral and dorsal halves, respectively. Segregation into a ventro-lateral forskolin-independent odorant processing stream and a medio-dorsal forskolin-dependent stream is visible. Below and side: Positions of responsive regions along the anterior-posterior and lateral-medial axes were measured and the probability density along those axes was estimated using Gaussian kernel density estimation. Regions responding to odorants but not to forskolin (fsk-) are shown in blue, regions responding either to odorants and forskolin or only forskolin in orange (fsk+). The 2D positional data of responsive regions of 11 vMOB hemispheres (n_PRO_ = 6, n_MET_ = 5) from 8 different animals (n_PRO_ = 4, n_MET_ = 4) was compared. First, between the ventral halves (below, left) and dorsal halves (below, right) of the vMOB as well as between prometamorphotic (side, above) and metamorphotic animals (side, below). White and black crosses indicate the peaks of the density estimates for the fsk-/fsk+ processing streams, respectively. Each contour-level corresponds to an iso-proportion of the density distributions (20% of the distribution for each level). The distributions in each plot were normalized to the number of responding regions. Margin plots show the positions of individual responsive regions and their estimated density distribution along the respective axis. The responding regions belonging to the two streams show different distributions in all four plots, with the odorant positive/forskolin negative group located laterally and posteriorly and forskolin-responsive regions located medially and anteriorly. **(b)** Changes of the fluorescence signal of responding regions upon odorant/forskolin stimulations over time in seconds are shown. Fluorescence changes of individual time traces are presented as percentages of the maximum fluorescence of this region (%). Black lines in the background are time traces of all individual responsive regions belonging to a specific tuning profile. Colored lines represent the average traces of each tuning profile. The six most frequent response profiles are shown. Regions are tuned to amino acids, amines, or bile acids individually or in different combinations and either responsive or non-responsive to forskolin. Stacked bar plots show the percentual share of individual tuning profiles among all identified responsive regions (n = 270) in the ventral and dorsal half of the vMOB in prometamorphotic and metamorphotic animals (ventral: n_Pro_ = 110, n_MET_ = 73; dorsal: n_Pro_ = 31, n_MET_ = 56). Blue shaded profiles are forskolin-non responsive (n = 111), yellow/orange profiles are forskolin-responsive (n = 159). A anterior, AA amino acids, AM amines, BA bile acids, FSK forskolin, L lateral, M medial, MET metamorphosis proper, ON olfactory nerve, P posterior, PRO prometamorphosis, vMOB ventral main olfactory bulb.

**Figure 4:**
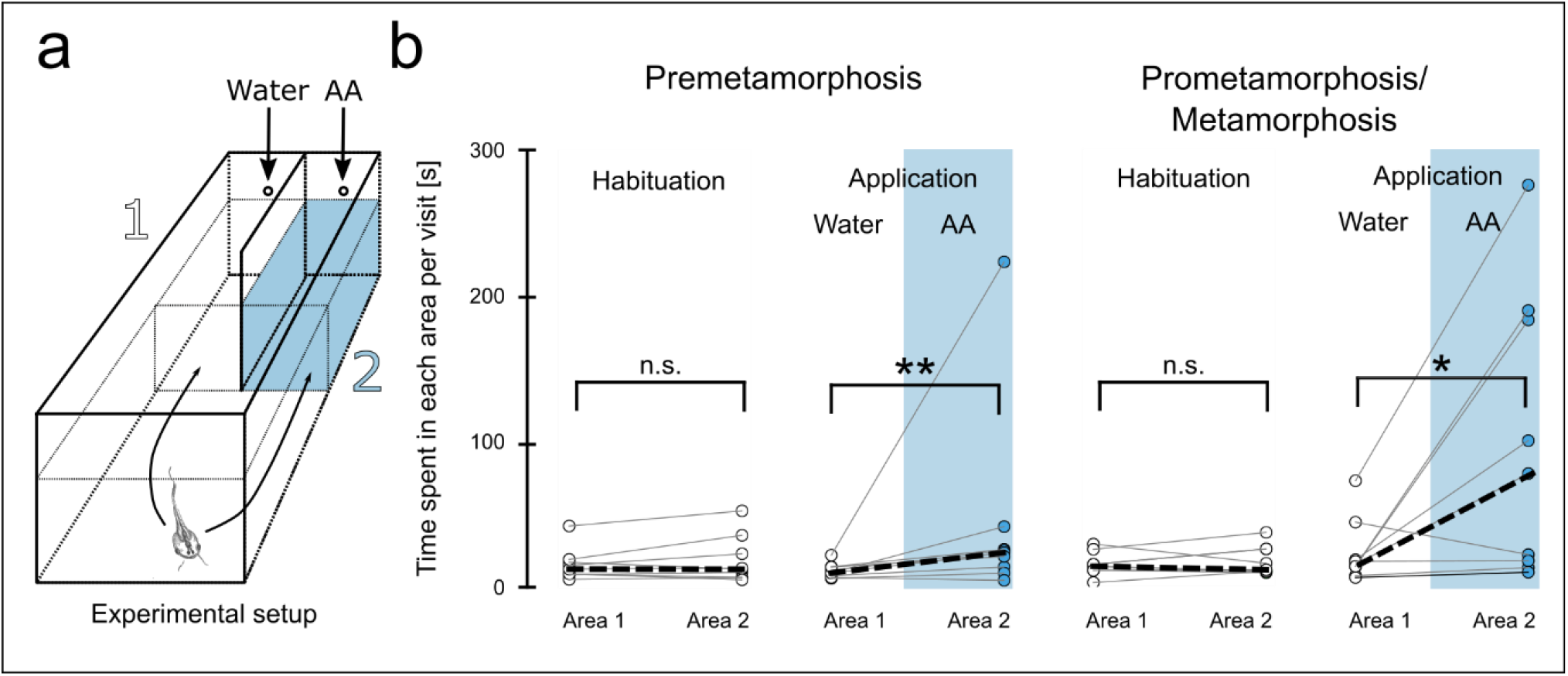
Behavioral responses to amino acids are stable until metamorphotic climax. **(a)** Experimental setup is shown. Tadpoles could freely move between area 1 and 2, which are separated by a dividing wall. After two hours of habituation, a water control stimulus was applied in area 1 and amino acids in area 2. **(b)** The average time each animal spent per visit to area 1 and area 2 is shown for the habituation period (left plot) and after stimulus application (right plot) for premetamorphotic animals (n = 9) and pro-/metamorphotic animals (n = 9). During the habituation period, no preference for either of the two areas was found. After stimulus application, both premetamorphotic animals, as well as animals of higher stages were found to spend significantly more time per visit in the amino acid area. *p < 0.05, **p < 0.01. AA amino acids, PRE premetamorphosis, PRO prometamorphosis, MET metamorphosis proper.

**Figure 5:**
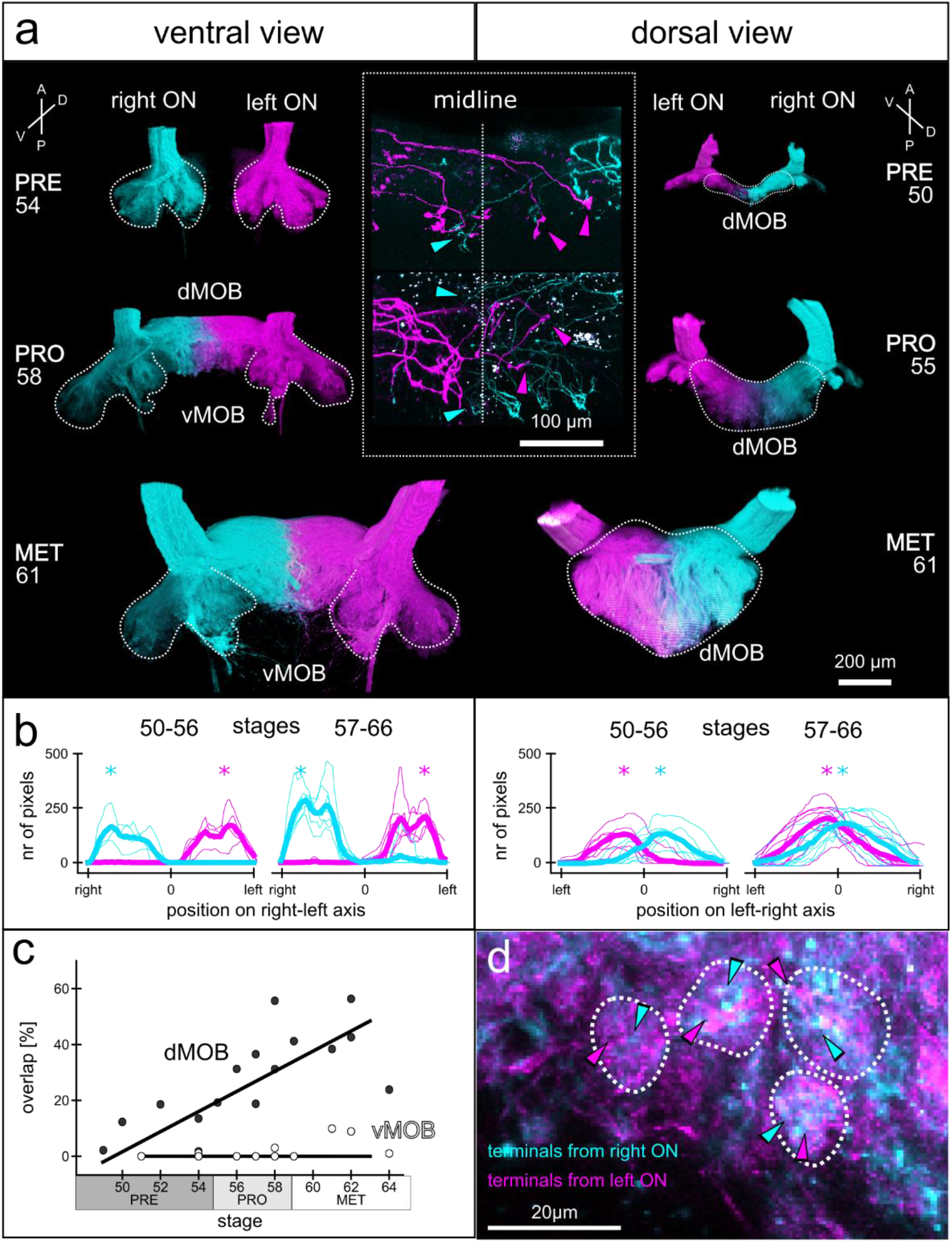
During metamorphosis, ORN axons from the left and right olfactory epithelia form a single dMOB projection field with increasing bilateral innervation overlap. **(a)** Images of ORN axonal projections in the vMOB (left images, ventral view) and the dMOB (right images, dorsal view) in a pre-, pro-, and metamorphotic animal. Projections from the left (magenta) and right (cyan) olfactory epithelia (both PC and MC) were traced with two different fluorophore-coupled dextran dyes (Alexa Fluor 488 and 594 dextrans) via electroporation. White dotted lines indicate the outlines of ORN axonal projections. While the left and right projections in the vMOB are clearly separated, the dMOB progressively fuses around the midline. The insert in the middle shows a close-up of sparsely labeled ORN axons crossing the midline (vertical dotted line) innervating contra- and ipsilateral glomeruli in the dMOB. **(b)** The number of pixels containing fluorescence signal from the ORN projections from the left (magenta) and right (cyan) were counted at each position along the left-right axis. The thin lines represent individual animals and the bold lines the mean of all animals belonging to the two presented developmental groups (stages 50–56 and 57–66). The x-axis is scaled from the outermost right edge to the outermost left edge of axonal projections in the vMOB and mirrored around the midline in the dMOB, with 0 being the interhemispheric midline. In the vMOB (left plots), pixels around the midline did not contain fluorescence signal from ORN axons of either of the two epithelia, while in the dMOB (right plots), pixels around the midline contained signal from axons from the left and right epithelia. The asterisks highlight the position with the highest count of pixels containing signal of axons from the left (magenta) or right (cyan). The maxima are close to the midline in the dMOB of both developmental groups. **(c)** The overlapping volume of axonal projections from the left and the right relative to the entire ORN axonal projection volume was calculated for tadpoles of different developmental stages and separately for the vMOB (white dots) and the dMOB (black dots). While the vMOB projections have no overlapping volume, the innervation overlap in the dMOB increases during metamorphosis. A RANSAC regression line was fitted through the data. **(d)** A close-up of the projections around the midline in the dMOB shows, that single glomerular structures get input from both left and right olfactory epithelia (indicated by the magenta and cyan arrowheads, respectively). A anterior, D dorsal, dMOB dorsal main olfactory bulb, MET metamorphosis proper, ON olfactory nerve, P posterior, PRE premetamorphosis, PRO prometamorphosis, V ventral, vMOB ventral main olfactory bulb.

We analyzed a total of 270 responding regions and examined their tuning profiles and locations in the vMOB (Figure 3a, b). Responding regions varied significantly in size (average 770 ± 1264 μm^2^). We measured the relative position along the medial-lateral and anterior-posterior axis. Based on these positions, we estimated the probability density function along the two spatial axes to compare between the spatial distribution of forskolin-responsive (Figure 3a; orange; fsk+) and forskolin-non-responsive regions (blue; fsk-). We estimated the 2D probability density separately for the ventral and dorsal halves of all vMOBs (including pro- and metamorphotic animals) to add the third spatial dimension to the analysis (density plots at the bottom of Figure 3a). Additionally, we compared the probability densities between prometamorphotic and metamorphotic samples (including both ventral and dorsal halves; plots at the right, Figure 3a) to monitor possible changes during metamorphotic remodeling.

The data showed that regions responding to forskolin and consequently using cAMP as second messenger (fsk+) are generally distributed more medially and anteriorly, with the area of highest estimated density visible in dark orange/brown. Forskolin-non-responsive regions (fsk-) are instead distributed more laterally and posteriorly (area of highest estimated density in dark blue). The peak of the density estimates for forskolin-responsive and forskolin-non-responsive regions are marked by black and white crosses, respectively. These general locations for the peak of the density distributions hold true for both the ventral and dorsal halves of the vMOBs (bottom plots, Figure 3a) and the pro- and metamorphotic animals (right plots, Figure 3a).

However, almost no forskolin-independent regions were found in the dorsal halves of the examined vMOBs, resulting in a very low density in the corresponding plot. Also, forskolin-responsive regions in the dorsal halves can be found in more lateral positions when compared to the ventral halves. In metamorphotic animals, the highest density of forskolin-responsive regions is shifted more anteriorly and the overall density of forskolin-independent regions is lower when compared to earlier developmental stages. The lateral and posterior position of these regions remains however unchanged when compared to the prometamorphotic group.

In addition to their reactiveness to forskolin (fsk+ and fsk-), we categorized the responsive regions based on their response-profiles to the three odorant mixtures (amino acids, bile acids, amines; Figure 3b). Of the 270 responsive regions, 111 did not respond to forskolin but showed odor-evoked responses to at least one of the three applied mixtures. 159 regions responded either to forskolin only or to the odorant mixtures in addition to forskolin. In total, we found 15 different response-combinations, seven of which were forskolin-independent and eight which showed responses upon forskolin application. The most frequent response profiles were amino acid-responsive, but not responsive to forskolin (bright blue trace, n = 77), followed by regions responding to forskolin only (bottom trace, n = 56) and forskolin-responsive regions also responsive to amines (second trace from the bottom, n = 30). Some regions showed responses to multiple of the applied stimuli (traces, Figure 3b).

We next examined the composition of responsive regions in both the ventral and dorsal part of the vMOB separately for pro- and metamorphotic animals (Figure 3b, stacked bar plots). Amino acid-responsive/forskolin-non-responsive regions were more frequent in the ventral part of the vMOBs of both prometamorphotic and metamorphotic animals (41% and 32%, respectively) than in the dorsal part of the vMOBs (13% and 9%). Contrarily, regions responsive to forskolin only are more frequent in the dorsal part of the vMOBs (prometamorphotic 26% and metamorphotic 40%) compared to the ventral halves of the vMOBs (9% and 22%). In total, regions also or only responding to forskolin make up 43% in the ventral parts of pro- and 55% in the ventral part of metamorphotic animals. In the dorsal part of the vMOBs, forskolin-responsive regions make up 87% (prometamorphotic animals) and 80% (metamorphotic animals).

Overall, these results strongly suggest that the segregation into two major odorant processing streams in the vMOB of tadpoles remains intact and unchanged even during the major metamorphotic rewiring processes. Since these experiments solely highlight odor representation on the glomerular level, we next wanted to examine odor responses on a behavioral level to ascertain their functional stability during metamorphosis.

### Tadpoles of different developmental stages show behavioral response to amino acids

We performed a behavioral choice assay using tadpoles of different metamorphotic stages up to metamorphotic climax to assess their behavioral response to amino acid stimulation in a choice tank (Figure 4a). Tadpoles were first left to swim freely for a two-hour habituation period, and the average time they spent per visit in the two areas left (area 1, Figure 4a) and right (area 2) of the dividing wall of the tank was recorded. During the two-hour habituation period, no stimulus was applied. Tadpoles showed no inherent preference to spend more time per visit in either side of the tank (premetamorphotic tadpoles: median 13 sec in area 1 and 13 sec in area 2; n = 9 animals; pro/metamorphotic tadpoles: median 14 sec in area 1 and 12 sec in area 2; n = 9; Figure 4b).

After habituation, tadpoles were simultaneously presented with water control and an amino acid application, and the time they spent per visit in the water area (area 1), and the amino acid area (area 2) was averaged over a 20-minute recording period (Figure 4b). Tadpoles preferred to spend more time per visit in the amino acid area over the water control area (premetamorphotic tadpoles: median 23 sec in the amino acid area, 10 sec in the control area, n = 9; prometamorphotic/metamorphotic tadpoles: median 78 sec in the amino acid area, 15 sec in the control area, n = 9; Figure 4b). Our results suggest that tadpoles are generally able to detect the presence of amino acids even during the substantial rewiring of the vMOB.

### Incoming ORN axons in the dMOB cross the midline and form a single projection area

While the neuronal network in the vMOB is rewired and ORN axons originating in the PC are replaced by new MC fibers, the dMOB network only starts to form during metamorphosis. We traced the ORN axons originating in the left and the right olfactory epithelium (both PC and MC) separately using two different fluorophore-coupled dextrans introduced by bulk electroporation and found several differences in the network structure between the vMOB and the dMOB (Figure 5). Figure 5a shows the 3D depictions of ORN axonal projections traced in a pre-, pro-, and metamorphotic animal. The images on the left and on the right show the same specimens in a ventral and dorsal view, respectively.

In the vMOB, the axons coming in from the left and right olfactory nerve project to two vMOB projection fields that are spatially separated at the interhemispheric midline (highlighted by the white dotted lines in the left images, Figure 5a). The general outline of the glomerular projections in the vMOB remains constant throughout metamorphosis and only slightly increases in overall volume. Contrastingly, the first axonal projections reach the dMOB around stage 50 (Figure 5a, top right) and project towards the interhemispheric midline. Until the onset of prometamorphosis, the incoming fibers from both olfactory nerves have formed a single projection area around the midline, which grows until the end of metamorphotic development. (Figure 5a, right). During prometamorphosis and metamorphosis proper, the hemispheres of the dMOB projections are not clearly separated anymore since axons cross the midline from both sides, frequently innervating glomeruli on the contralateral side. This feature of the dMOB becomes even more apparent when looking at sparse cell labeling (inset, Figure 5a). Among the ORN axons crossing to the contralateral side, we found axons only connecting to the ipsilateral or contralateral side, but also some axons bifurcating and innervating glomeruli on both sides of the midline.

To quantify the positions of incoming ORN axons from the left (magenta, Figure 5) and right epithelia (cyan), we counted pixels containing fluorescence signal from the ORN axons of the two epithelia along the left-right axis of the vMOB (left panel , Figure 5b) and dMOB (right panel, Figure 5b). The number of pixels containing signal from axons originating in the left or right epithelium is plotted as a function of distance from the interhemispheric midline (0) to the outermost left and right edges of the axonal projections (Figure 5b). In the vMOB (left panel, Figure 5b), the curves have symmetrical peaks on both sides of the interhemispheric midline (at approx. 80% of the distance to the edges; asterisks, Figure 5b), while in immediate proximity to the midline, no pixels containing fluorescence signal could be counted. This pattern did not change from earlier stages (50–56; n = 4 animals) to later stages (57–66; n = 7) during metamorphosis. In the dMOB, on the other hand (right panel, Figure 5b), the symmetrical peaks were located closer to the midline, at approx. 15% (asterisks) of the distance to the edges in animal of stages 50–56 (n = 7) with some overlap around the midline (area under both curves). In the later metamorphotic stages (57–66; n = 9), the two peaks are almost on the midline (5% of the distance to the edges; asterisks), with even more substantial overlap.

We further quantified the percentual share of innervation overlap relative to the total projections in animals of different stages for the vMOB (Figure 5c, white dots) and dMOB (Figure 5c, black dots). Overlap was computed by counting the number of pixels containing fluorescence signal from both epithelia (magenta and cyan, Figure 5) relative to the total number of pixels containing any fluorescence signal. The average overlap of the right and left ORN axons in the vMOB amounts to 2.2 ± 3.7% (animals of all examined stages; n = 11; Figure 5c).

The percentage of overlap in the dMOB increases linearly throughout metamorphosis (premetamorphosis: 10 ± 7%, n = 5; prometamorphosis: 32 ± 14%, n = 6; metamorphosis proper: 40 ± 12%, n = 5; Figure 5c). In addition, our results show that a single glomerular structure in the dMOB can be composed of axon terminals from both sides (magenta and cyan arrowheads, Figure 5d). Since glomeruli are the sites of synaptic connection to postsynaptic bulbar neurons, the left-right overlap could be the basis of a bilateral signal integration system of the dMOB.

### The populations of projection neurons in the vMOB and the dMOB are distinct

The connection between peripheral receptor neurons and the glomeruli differs between the vMOB and the dMOB. To understand, how these projections from the periphery connect to the postsynaptic projection neurons, we labelled and reconstructed single projection neurons in the vMOB and dMOB of postmetamorphotic *Xenopus laevis* (stage 66) (Figure 6).

**Figure 6:**
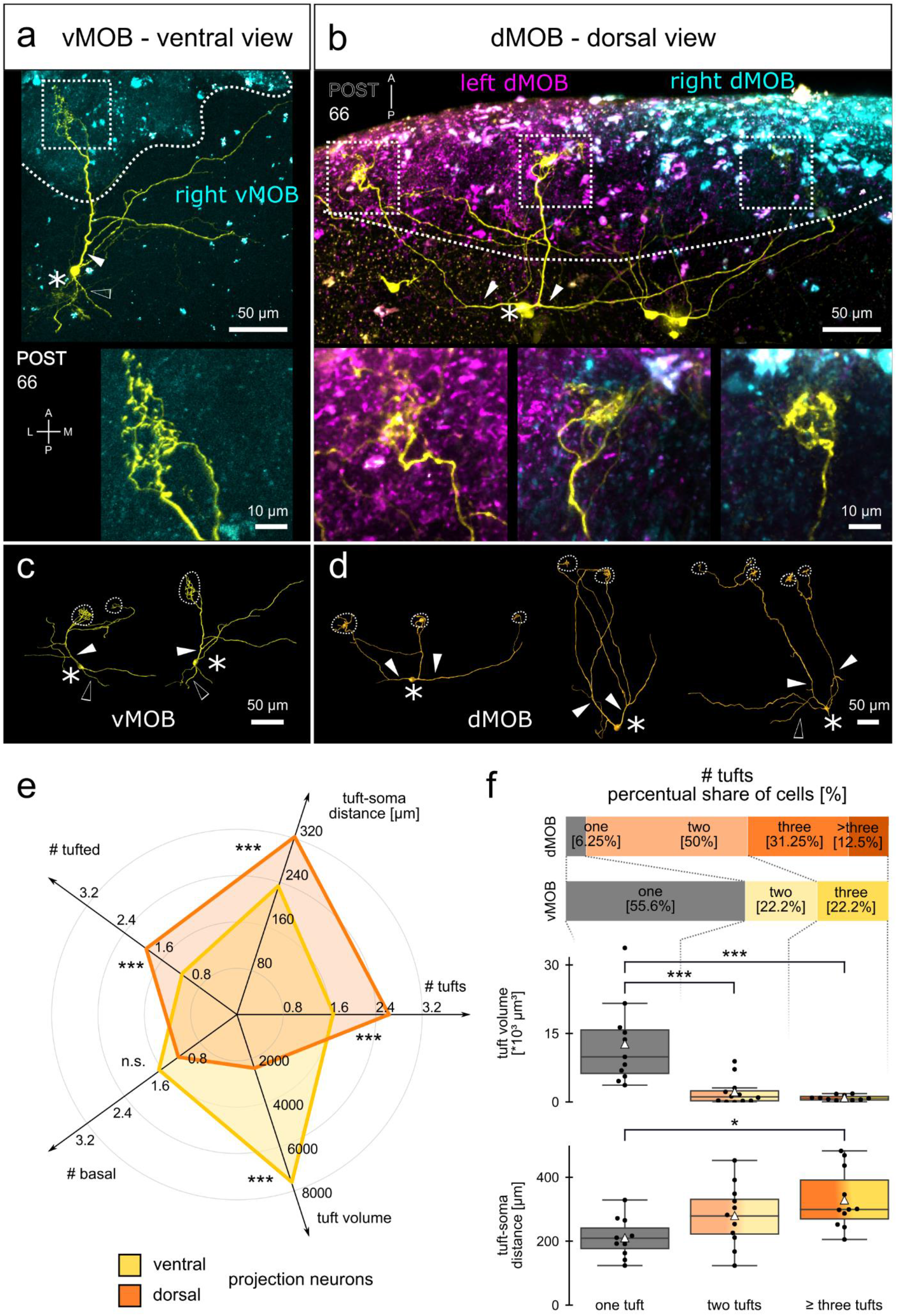
Different morphology of projection neurons in the vMOB and dMOB of postmetamorphotic *Xenopus*. ORN axonal projections from the olfactory epithelia (both PC and MC) were traced via electroporation of fluorophore-coupled dextrans (left: Cascade Blue dextran, magenta; right: Alexa Fluor 594 dextran, cyan), and projection neurons (yellow) with Alexa Fluor 488 dextran via sparse cell electroporation in the vMOB **(a)** and dMOB **(b)**. White dotted lines indicate the ORN projections, tufts (white dotted squares) are shown in a higher magnification below. Reconstructions of representative neurons in the vMOB **(c)** and dMOB **(d)** are shown. Asterisks indicate the projection neuron somata, filled white arrowheads primary tufted dendritic branches and empty arrowheads primary basal neurites without tufted terminals. **(e)** Each radial axis on the radar chart represents a morphological descriptor of the projection neurons. The means of the descriptors of projection neurons in the vMOB (yellow; n = 18) and the dMOB (orange; n = 16) are shown. Dorsal projection neurons have a higher number of tufts, a longer average distance between soma and tufts and more primary tufted dendrites. Neurons in the vMOB have a significantly higher tuft volume. The number of primary basal neurites is similar between the groups. **(f)** The stacked barplots depict the percentual share of uni-tufted (grey) and multi-tufted projection neurons (yellow, vMOB; orange, dMOB). A higher number of uni-tufted projection neurons was found in the vMOB (>50%), while >90% of neurons in the dMOB terminated in at least two tufts, maximally in five tufts. Boxplots: Average tuft volumes (above) and average distance between tufts and somata (below) are compared between uni-tufted neurons (grey, left, n = 11), bi-tufted neurons (yellow/orange, middle, n = 12) and neurons with three or more tufts (dark yellow/dark orange, right, n = 11). White triangles depict the means. Tuft volume decreases with a higher number of tufts, while tuft-soma distance increases. The groups contain neurons from both the vMOB and the dMOB. *p < 0.05, **p < 0.01, ***p < 0.005. A anterior, dMOB dorsal main olfactory bulb, L lateral, M medial, ON olfactory nerve, P posterior, POST postmetamorphosis, vMOB ventral main olfactory bulb, PRE premetamorphosis, PRO prometamorphosis, MET metamorphosis proper, vMOB ventral main olfactory bulb.

Projection neurons in the vMOB (n = 18 neurons, Figure 6a, c) and dMOB (n = 16, Figure 6b, d) share some general morphological features. Both groups have one or multiple primary dendrites (filled arrowheads in Figure 6c, d) originating from the soma (white asterisks in Figure 6) and terminating in highly branched dendritic tufts connecting to one or multiple glomeruli (insets in Figure 6a, b). A representative projection neuron with a single tuft in the right vMOB is shown in Figure 6a, and a multi-tufted neuron in the dMOB in Figure 6b. While the tufts of neurons in the vMOB get synaptic input only from ipsilateral ORN projections (cyan projections in Figure 6a), multiple tufts of a single projection neuron in the dMOB often project to both hemispheres, possibly integrating synaptic ipsilateral and contralateral ORN input (cyan and magenta in Figure 6b). In addition to the dendrites receiving input from the dendritic tufts, most cells also have secondary dendritic branches with blunt endings and basal neurites (empty arrowheads, Figure 6). Figures 6c and 6d show reconstructions of representative projection neurons in the vMOB and dMOB that connect to a different number of glomeruli (white dotted circles).

From the reconstructions, we extracted several descriptors for the projection neurons (Supplementary video 1) and compared the measurements of neurons labeled in the vMOB and the dMOB (Figure 6e). The tuft-soma distance along the dendrites (for cells with multiple tufts, the distance was averaged) was significantly longer in the dMOB cells (323 ± 96 μm) compared to the vMOB cells (230 ± 70 μm; p = 0.0046; Figure 6e). Projection neurons in the vMOB and dMOB connected to an average of 1.7 ± 0.8 and 2.6 ± 1.1 dendritic tufts (p = 0.043), respectively, with an average tuft volume of 7679 ± 8208 μm^3^ and 2435 ± 5410 μm^3^ (p = 0.0003). We classified neurite branches along two axes into primary (originating in the soma) or secondary (branching off from primary or other secondary branches; Supplementary video 1) as well as dendritic/tufted (connected to the tufts) or basal neurites (not connected to the tufts). Neurons in the vMOB had 1.2 ± 0.5 primary-tufted dendrites and 1.7 ± 1 primary-basal neurites. The dMOB neurons exhibited significantly more of the former (1.9 ± 0.7; p = 0.0003) and a similar number of the latter (1.3 ± 1.4; Figure 6e). The count of secondary-tufted dendrites was similar among the two groups (vMOB 3.9 ± 2.1, dMOB 3.8 ± 3.4) while the dMOB cells had more secondary-basal branches (7.1 ± 2.7) compared to the vMOB (4.4 ± 2.7; p = 0.004).

The most apparent difference between the two neuronal populations is the share of uni- and multi-tufted cell morphologies (Figure 6f, barplots). 55.6% of neurons labeled in the vMOB are uni-tufted (grey, Figure 6f), while only a single uni-tufted cell has been found in the dMOB (6.25%). The bi- and tri-tufted neurons both make up 22.2% of the vMOB population (yellow in Figure 6f). In the dMOB, the projection neurons maximally had five tufts. The biggest share of the population was bi-tufted and tri-tufted cells with 50% and 31.25%, respectively (orange in Figure 6f).

In the next step of the analysis, we pooled the data based on the number of tufts (one, n = 11; two, n = 12; three or more, n = 11; boxplots Figure 6f), combining both neurons from the vMOB and the dMOB. Uni-tufted neurons had the biggest tuft volume (12711 ± 8921 μm^3^; Figure 6f, middle), significantly bigger than the average tuft volume of the multi-tufted cells (bi-tufted: 2246 ± 2911 μm^3^, p = 0.001; three or more tufts: 946 ± 600 μm^3^, p = 0.0002). Inversely, the mean dendritic distance between tuft and soma increased with a higher number of tufts, measuring 211 ± 60 μm in uni-tufted, 281 ± 93 μm in bi-tufted, and 330 ± 93 μm in cells with three or more tufts (Figure 6f, below).

As an additional descriptor for all multi-tufted cells, we measured the average spatial distance between two tufts of a cell as an estimate of spatial span of the neuron. Tufts of bi-tufted neurons were on average 146 ± 93 μm apart, with a maximum distance of 325 μm, while we measured 128 ± 74 μm for neurons with three or more tufts. Comparing all multi-tufted projection neurons in the vMOB (n = 8) and dMOB (n = 15) showed that the inter-tuft distance is significantly bigger in the dMOB (178 ± 73 μm; p = 0.0003) compared to the vMOB (62 ± 32 μm). Inversely, the volume of multi-tufted neurons in the vMOB was bigger than in the dMOB (2500 ± 2671 μm^3^; 1157 ± 1836 μm^3^; p = 0.008).

In summary, the projection neurons in the dMOB develop multiple small dendritic tufts, which receive input from spatially distant glomeruli, while at least half of the vMOB neurons have a single bigger tuft receiving input from only one glomerulus. These characteristics further support the idea that the network and processing logic of the vMOB and the dMOB are distinct. It remains to be elucidated how this links to their respective functions in the detection of water- and airborne odorants, respectively.

## Discussion

### ORN projections from the PC and MC form two distinct subsystems during metamorphosis

During metamorphosis, the main olfactory system of most amphibians must undergo a complete transformation to adapt to the terrestrial lifestyle of the adult frog. In the case of *Xenopus*, the aquatic main olfactory epithelium of the tadpole transforms into a bi-modal system consisting of the MC epithelium, associated with detection of waterborne odorants, and the PC epithelium dedicated to aerial olfaction. This is supported by evidence showing that the larval PC and the adult MC both possess ciliated and microvillous ORNs (Hansen et al., 1998) and a similar set of olfactory receptors expressed (Amano and Gascuel, 2012; Syed et al., 2013, 2017) that are tuned to detect waterborne odorants like amino acids (Syed et al., 2017). The postmetamorphotic PC, on the other hand, putatively expresses receptor genes more closely related to the mammalian receptors responsive to volatile odors (Freitag et al., 1995, 1998).

The projections of the ORNs in adult *Xenopus* towards their glomerular targets in the MOB have been described based on their lectin binding pattern (Key and Giorgi, 1986; Hofmann and Meyer, 1991; Franceschini et al., 1992). Fibers coming from the adult MC were found to be soybean-agglutinin positive and innervate the ventrolateral MOB, while the ‘aerial-fibers’ from the PC were soybean-agglutinin negative and projected into the dorsomedial MOB (Hofmann and Meyer, 1991; Gaudin and Gascuel, 2005). In a thorough study of ORN projection fields in the MOB during development and metamorphosis, Gaudin and Gascuel describe that the dMOB (called PF9 in their study) increases significantly in size (28 times increase) between stage 50 and 59, while the increase factor dropped to 1.4 in stages between 59 and 64 (Gaudin and Gascuel, 2005). Our results confirm these findings (Figure 1). We did not evaluate the absolute growth of the olfactory bulb structures but the changes in the percentual share of the projection zones in the vMOB and the dMOB. We found that the biggest increase in the relative size of the dMOB happened until stages 55/56 with an increase from 0% to around 60% of the total volume, while the increase was only 10% from stages 57 to 64 (Figure 1c). We additionally found that around stage 54, the projection fields of the dMOB and the vMOB already have approximately the same volume.

We investigated incoming fibers to the dMOB in animals throughout metamorphosis and found that input into the dMOB was solely originating from the PC. This is in accordance with prior studies (Reiss and Burd, 1997b; Burd, 1999). We did not find any fibers from the MC (Figure 2c, right bar plot). The vMOB of the tadpoles is innervated by ORN axons from the PC. Around stage 52, the first axons from the newly formed MC reach the vMOB glomeruli (Reiss and Burd, 1997b; Burd, 1999). Reiss and Burd observed that from stages 52 to 58, PC axons could still be observed in the vMOB, while after that stage, the PC afferents have completely vanished from the vMOB, leaving only the MC axon terminals (Reiss and Burd, 1997b).

We have quantified this dynamic shift in innervation in the vMOB and found that the relative innervation of the PC decreases gradually from stage 52 to around stage 61 (Figure 2c, left plot). We still observed some PC axons after the onset of metamorphotic climax (stage 58), even if by that time in the development, vMOB glomeruli are already majorly innervated by incoming MC ORN axons. Our data also is in accordance with results showing a peak in cellular apoptosis in the PC around stage 58 and later again around 62 (Dittrich et al., 2016). Some ORNs in the PC projecting to the vMOB could still be present at stages 58–62, undergoing apoptosis during the second apoptotic peak around stage 62, later than proposed by Reiss and Burd (Reiss and Burd, 1997b).

### Dynamic innervation shift of the vMOB does not disrupt glomerular or behavioral responses to odorants

During the described rewiring process in the vMOB, it is unclear so far whether the glomeruli retain their larval functionality, if the odorant response pattern is reorganized or if they temporarily lose a clear functional pattern. We imaged odorant induced response profiles of glomeruli in tadpoles up to stage 61 in the ‘water bulb’ of *Xenopus laevis* and found that the extensive fiber replacement shows little effect on the coarse spatial organization of odorant responses in the vMOB. Odorant mediated responses can be recorded in the glomeruli of the vMOB up until the metamorphotic climax, and their location within the glomerular cluster does not change significantly (Figure 3). Our data suggests that replacement of ORN projections from the larval PC by new ORN projections originating in the MC happens gradually, without interrupting the spatial configuration and responsiveness of glomeruli in the vMOB. Similarly, behavioral responses to amino acids persisted at least until the metamorphotic climax (Figure 4). This implies an intact functional connection between the vMOB and higher downstream brain centers.

A recent study in the semiaquatic bullfrog *Lithobates catesbeiana* using electro-olfactography (EOG) showed a reduced electrophysiological response to spirulina extract – a food stimulus – as well as to a single amino acid (L-alanine) in metamorphotic animals. Similarly, the behavioral preference for spirulina decreased during the metamorphotic climax (Heerema et al., 2020). The authors of this study hypothesized that this decrease could mirror the temporary cessation of feeding behavior displayed by many anuran tadpoles during the metamorphotic climax (Hourdry et al., 1996; Heerema et al., 2020). Alternatively, the reduced response to spirulina could stem from an ontogenetic shift to a terrestrial lifestyle in the bullfrog and a decrease in responsiveness to aquatic cues such as spirulina.

Even though we observed less odorant responsive regions in the metamorphotic group when compared to the prometamorphotic group, we did not see a complete cessation of olfactory responsiveness up to animals of stage 61. Also, metamorphotic animals still showed behavioral responses to amino acids. Since *Xenopus* remains fully aquatic after metamorphosis is completed, it is conceivable, that there is less or no reduction of functionality in the water-smelling system during metamorphotic climax than in species that change their lifestyle. This idea is further supported by a recent study showing that the glomerular clusters in the water bulb of *Dendrobates tinctorius,* a terrestrial frog species, seem gradually reduced until postmetamorphosis, when compared to the vMOB in *Xenopus* (Weiss et al., 2020a).

In tadpoles of *Xenopus laevis*, the spatial organization of odorant responses in the main olfactory epithelium and the MOB have been extensively studied (Manzini and Schild, 2003, 2010; Gliem et al., 2013; Syed et al., 2017), but little to no functional imaging data exists for the metamorphotic and postmetamorphotic stages. In a comprehensive study from the epithelium to the MOB of tadpoles, it was shown that the MOB projections can be subdivided into two parallel processing streams (Gliem et al., 2013). Putatively microvillous ORNs projecting to the lateral glomeruli have a cAMP-independent second messenger pathway (Manzini et al., 2002; Manzini and Schild, 2003) and are mostly responsive to amino acids (Gliem et al., 2013). The medial stream relies on the canonical cAMP-dependent pathway instead and is most probably formed by ciliated ORNs (Gliem et al., 2013). It was hypothesized that the lateral stream could be linked to ORNs expressing vomeronasal type 2 olfactory receptors (V2Rs) expressed in the lateral main olfactory epithelium of the larvae (Gliem et al., 2013; Syed et al., 2013, 2017). The expression pattern of the V2Rs and the response pattern to amino acids gradually shifts to the adult MC during metamorphosis (Syed et al., 2017).

We show that the segregation between the lateral forskolin-non-responsive/ cAMP-independent pathway and the forskolin-responsive/cAMP-dependent medial pathway is retained until stage 61 tadpoles (Figure 3). The probability density distributions of responsive regions in Figure 3a show that forskolin-induced calcium transients (orange) are missing in the ventrolateral glomerular cluster. Glomeruli in this part of the vMOB are instead responsive to odorant stimulation, particularly to stimulation with amino acids (blue). Our results also show that the cAMP-independent pathway is restricted to the ventral parts of the vMOB, while dorsal glomeruli are forskolin-responsive/cAMP-dependent. It is to note that many responsive regions analyzed in this study only responded to forskolin, but not to any of the applied odorants. Based on previous experiments in *Xenopus* larvae (Gliem et al., 2013), we assume that these regions are responsive to different odorant groups, which were not individually applied in the present study. We also observed forskolin-induced responses in dMOB glomeruli in some of the animals, which complies with the idea of exclusively ciliated ORNs using the cAMP pathway projecting to this region of the MOB (Hansen et al., 1998). Future functional experiments using airborne odorants will have to examine the odor tunings of these ORNs.

Odorant receptor molecules themselves are thought to be a main determinant factor for ORN axonal targeting in the MOB in vertebrates (Feinstein and Mombaerts, 2004; Mombaerts, 2006). Since the MC expresses the same olfactory receptors as the larval PC epithelium (Amano and Gascuel, 2012; Syed et al., 2017), it seems likely that the vMOB odor map also remains constant. Axon targeting mechanisms in the vMOB of *Xenopus* are likely to differ from the targeting in the dMOB since it relies on different odorant receptor families. Additionally, axon targeting does not seem to be restricted to the ipsilateral hemisphere since ORN fibers cross extensively over the midline. Up to date, no odorant map of the dorsal projections is available. To our knowledge, it is also unclear whether the dMOB of anurans is functionally organized symmetrically or whether ORN axons expressing a given receptor in the left and right PC project their axons to a single glomerulus.

### The fusion of the olfactory bulb is not exclusive to anurans

The fused dMOB is not a feature exclusive to the anurans. It has also been described in some fishes and bird species (Nieuwenhuys, 1966). It has been studied in more detail in passeriform birds (Huber and Crosby, 1929; Yokosuka et al., 2009b, 2009a; Corfield et al., 2015). However, in birds, the fusion on the gross anatomical level does not seem to relate to a functional overlap. In the Japanese Jungle Crow, it was shown that even though the olfactory bulb appears as one single mass, the olfactory nerves project separately to the left and right glomerular layer, with no significant left-right overlap (Yokosuka et al., 2009b). In contrast, the morphological results in the present study (Figure 5), as well as electrophysiological studies in ranid frogs (Jiang and Holley, 1992b), show a significant overlap between the ORN axon projections.

It is generally assumed that the relative size of the olfactory bulbs is correlated with the importance of olfactory capability across vertebrates (Corfield et al., 2015; Yopak et al., 2015). Among birds, the Passeriformes only possess a very small olfactory bulb in relation to the whole brain size (Corfield et al., 2015), suggesting a relative loss of reliance on their sense of smell. This is partially explained by the fact that these birds, especially the corvids, have evolved higher cognitive functions (Emery, 2006; Corfield et al., 2015). Their small, fused olfactory bulb could thus be a sign of evolutionary degeneration of olfactory function. In anurans, the fused olfactory bulb has been described in different species (Hoffman, 1963; Ebbesson et al., 1986; Scalia et al., 1991; Jiang and Holley, 1992b), independently of their habitat and the degrees to which they rely on olfaction. It seems thus unlikely that the fusion of the olfactory bulb can be interpreted as a sign of olfactory degeneration in the anurans as well.

The evolutionary absence of the fused olfactory bulb in most vertebrates suggests that it is a derived evolutionary feature in both birds and frogs rather than an ancestral trait. Also, the close sister clade of the anurans, the salamanders, have two separated olfactory bulbs (Eisthen and Polese, 2007). We thus propose a different origin of this feature and a putatively different function in the various animal groups, which remains to be explored.

### Projection neurons in the vMOB and the dMOB are morphologically different

The morphology of the primary projection neurons in the olfactory bulb has been extensively studied in a variety of vertebrates (for review, see Dryer and Graziadei, 1994; Nieuwenhuys, 1966). Mammalian projection neurons are divided into mitral and tufted cells, with further subtypes being distinguished (Nagayama et al., 2014). Mitral cell somata are generally bigger in size and located more caudally, tufted cells are smaller and their somata are located closer to the glomerular layer (Mori et al., 1983; Macrides et al., 1985; Dryer and Graziadei, 1994; Nagayama et al., 2014). In amphibians, it is still debated whether the population of projection neurons in the MOB can be subdivided (Herrick, 1924; Scalia et al., 1991; Jiang and Holley, 1992a; Laberge, 2008). Morphological (Scalia et al., 1991; Jiang and Holley, 1992a) and electrophysiological studies (Jiang and Holley, 1992b) in adult ranid frogs propose a similar subdivision into tufted-like and mitral-like projection neurons in anurans as well.

In the present study, we tackled the question of whether the projection neuron populations in the vMOB (water system) and the dMOB (air system) of the aquatic *Xenopus* are morphologically different. It is described for amphibians, reptiles, and fishes (Dryer and Graziadei, 1994; Imamura et al., 2020), that projection neurons possess multiple primary dendrites that terminate in glomerular tufts and a variable number of secondary dendrites without tufts (Nieuwenhuys, 1966; Scalia et al., 1991; Dryer and Graziadei, 1994). We found that the number of secondary dendrites was comparable for cells in the vMOB (3.9 ± 2.1) and the dMOB (3.8 ± 3.4), which also is comparable to quantifications in ranid frogs (between 1–6) (Jiang and Holley, 1992a), salamanders (0–5; Laberge, 2008), rabbits (2–5; Mori et al., 1983), rats (2–9; Dryer and Graziadei, 1994; Orona et al., 1984) and hamsters (1.7–4.7; Macrides and Schneider, 1982). Interestingly, the cells in the dMOB had a higher number of primary dendrites originating in the soma than vMOB cells, which mostly exhibit a single primary dendrite. This could enable dMOB cells to integrate independent dendritic input in the cell soma, while vMOB somata only receive a single dendritic input.

While the single primary dendrite of mammalian mitral/tufted cells in the MOB connect to a single glomerulus, projection neurons of earlier diverging vertebrates often innervate multiple glomeruli (Nieuwenhuys, 1966; Dryer and Graziadei, 1994; Nezlin et al., 2003). In the population analysis of projection neurons in ranid frogs, around 50% of examined neurons were found to be connected to a single glomerulus, ~30% were bi-glomerular, and 20% innervated more than two glomeruli (Jiang and Holley, 1992a). This distribution is quite close to the counts we obtained in the vMOB (56% uni-, 22% bi-, 22% tri-glomerular: Figure 6f). Interestingly, the number of glomeruli innervated by a single ORN axon in anurans also follows a similar distribution across different species and developmental stages (tadpoles 41% uni-glomerular, 59% multi-glomerular; juveniles: 50% uni, 50% multi-glomerular; Weiss et al., 2020). It is still unclear if there could be a functional correlation between the different types of these distributions.

The projection neuron population in the dMOB is almost exclusively multi-tufted, with very small tuft volumes and longer distances between somata and tufts as well as between tufts (Figure 6f). Generally, we noted that tuft volumes were bigger in uni-tufted-cells compared to multi-tufted cells, while distances were shorter in uni-than multi-tufted cells. A recent study in zebrafish suggested that the presence of uni- and multi-glomerular projection neurons hint towards the coexistence of distinct odor processing logics (Braubach and Croll, 2021). The authors hypothesize that uni-glomerular projection neurons are primarily associated with glomeruli narrowly tuned to specific odors, eliciting specific behavioral responses (Yabuki et al., 2016; Dieris et al., 2017; Wakisaka et al., 2017). Multi-glomerular neurons, on the other hand, are associated with smaller glomeruli, possibly implicated in integrative odor processing of stimuli like bile acids or amino acids (Braubach and Croll, 2021). Interestingly, a similar relationship between smaller glomeruli and multi-glomerular connectivity can also be observed in the completely unrelated locust olfactory system (Ignell et al., 2001).

Based on our data, we propose that the vMOB of *Xenopus* displays a co-existence of multiple wiring logics based on uni- and multi-tufted projection neurons, similar to what was shown in zebrafish (Braubach and Croll, 2021). In contrast, the dMOB seems to be primarily associated with multi-glomerular projection neurons possibly involved in integrative odor coding. In the light of the zebrafish data presented above, *Xenopus* could have some hardwired behavioral responses based on specific aquatic cues processed via the uni-glomerular system in the water bulb, while volatile odors could generally be processed in a combinatorial way.

Additionally, the fact that the dMOB is a single unpaired structure and projection neurons connect to glomeruli in both hemispheres suggests that this system also integrates information from the left and the right olfactory epithelia. This seems to add another level of information integration to the detection of volatile odorants in the frogs.

### Putative ecological and behavioral relevance of the dMOB

The presence of a well-developed system to sample airborne odorants seems counterintuitive in a fully aquatic frog. It has, however, been reported that *Xenopus* move overland mostly in search of other water bodies or food (Du Plessis, 1966; Measey, 2016). It has also been suggested that olfactory cues might play a role in overland orientation and migrations (Savage, 1965). In the present study, we show how the neuronal circuit of the dorsal ‘air-bulb’ in *Xenopu*s differs from the ventral ‘water bulb’ and most other vertebrate MOBs. The bilateral innervation of both ORNs and projection neurons gives it a high degree of integrative power. The importance of integration of bilateral sensory information has been shown for other sensory systems, but its role in olfaction has only been shown in rare instances (Catania, 2013). Among anurans, the fusion of the two MOB hemispheres seems to be an ancestral trait with so far unknown function (Ebbesson et al., 1986; Scalia et al., 1991; Eisthen and Polese, 2007). The anuran dMOB could be a primary olfactory center integrating odorant information from the left and right nose in individual, bilaterally innervated glomeruli or by multiple tufts of individual projection neurons. This could e.g. facilitate the overland spatial orientation of *Xenopus*. Odorant-guided orientation and homing behavior is widespread in anurans. However little is known about the neuronal structures involved (reviewed in Weiss et al., 2021). It is intriguing to think that the special input integration in the dMOB could be part of a spatial integration center used in orientation behavior.

## Acknowledgments

We thank the members of the Manzini Lab for fruitful discussion.

## Author Contributions

L.W., T.H., and I.M. conceptualized the study. L.W., P.S.A, S.J.H. investigation; L.W., T.O., S.J.H. formal analysis, visualization L.W. writing of the original draft; L.W., P.S.A, T.O., S.J.H., T.H., and I.M. writing—review and editing the manuscript; T.H. and I.M. funding acquisition and resources of the manuscript; supervision of the article.

## Ethics statement

All experiments performed followed the guidelines of Laboratory Animal Research of the Institutional Care and Use Committee of the Justus-Liebig-University Giessen (V 54 – 19 c 20 15 h 01 GI 15/7 Nr. G 2/2019; 649_M; V 54 – 19 c 20 15 h 02 GI 15/7 kTV 7/2018).

## References

Amano, T., and Gascuel, J. (2012). Expression of odorant receptor family, type 2 OR in the aquatic olfactory cavity of amphibian frog Xenopus tropicalis. PLoS One 7, e33922. doi:10.1371/journal.pone.0033922.

Benzekri, N. A., and Reiss, J. O. (2012). Olfactory metamorphosis in the coastal tailed frog Ascaphus truei (Amphibia, Anura, Leiopelmatidae). J. Morphol. 273, 68–87. doi:10.1002/jmor.11008.

Braubach, O., and Croll, R. P. (2021). The glomerular network of the zebrafish olfactory bulb. Cell Tissue Res. 383, 255–271. doi:10.1007/s00441-020-03394-4.

Burd, G. D. (1999). “Development of the olfactory system in the African Clawed Frog, Xenopus Laevis,” in The Biology of Early Influences, eds. R. Hyson and F. Johnson (Boston, MA: Springer US), 153–170. doi:10.1007/978-0-585-29598-5_9.

Catania, K. C. (2013). Stereo and serial sniffing guide navigation to an odour source in a mammal. Nat. Commun. 4, 1441–1448. doi:10.1038/ncomms2444.

Corfield, J. R., Price, K., Iwaniuk, A. N., Gutiérrez-Ibáñez, C., Birkhead, T., and Wylie, D. R. (2015). Diversity in olfactory bulb size in birds reflects allometry, ecology, and phylogeny. Front. Neuroanat. 9, 102. doi:10.3389/fnana.2015.00102.

Dieris, M., Ahuja, G., Krishna, V., and Korsching, S. I. (2017). A single identified glomerulus in the zebrafish olfactory bulb carries the high-affinity response to death-associated odor cadaverine. Sci. Rep. 7, 40892. doi:10.1038/srep40892.

Dittrich, K., Kuttler, J., Hassenklöver, T., and Manzini, I. (2016). Metamorphic remodeling of the olfactory organ of the African clawed frog, Xenopus laevis. J. Comp. Neurol. 524, 986–998. doi:10.1002/cne.23887.

Dryer, L., and Graziadei, P. P. C. (1994). Mitral cell dendrites: a comparative approach. Anat. Embryol. (Berl). 189, 91–106. doi:10.1007/BF00185769.

Du Plessis, S. S. (1966). Stimulation of spawning in Xenopus laevis by fowl manure. Nature 211, 1092–1092. doi:10.1038/2111092a0.

Duellman, W. E., and Trueb, L. (1994). Biology of amphibians. Baltimore: John Hopkins University Press doi:10.2307/1445022.

Ebbesson, S. O. E., Bazer, G. T., and Jane, J. A. (1986). Some primary olfactory axons project to the contralateral olfactory bulb in Xenopus laevis. Neurosci. Lett. 65, 234–238. doi:10.1016/0304-3940(86)90311-3.

Eilers, P. H. C., and Boelens, H. F. M. (2005). Baseline correction with asymmetric least squares smoothing. Leiden Univ. Med. Cent. Rep. 1, 5.

Eisthen, H. L., and Polese, G. (2007). “Evolution of vertebrate olfactory subsystems,” in Evolution of Nervous Systems, Vol 2: Non-mammalian Vertebrates, eds. J. H. Kaas and T. H. Bullock (Oxford: Academic Press), 355–406.

Emery, N. J. (2006). Cognitive ornithology: The evolution of avian intelligence. Philos. Trans. R. Soc. B Biol. Sci. 361, 23–43. doi:10.1098/rstb.2005.1736.

Feinstein, P., and Mombaerts, P. (2004). A contextual model for axonal sorting into glomeruli in the mouse olfactory system. Cell 117, 817–831. doi:10.1016/j.cell.2004.05.011.

Föske, H. (1934). Das Geruchsorgan von Xenopus laevis. Z. Anat. Entwicklungsgesch. 103, 519–550. doi:10.1007/BF02118933.

Franceschini, V., Giorgi, P. P., and Ciani, F. (1992). Primary olfactory terminations in the forebrain of amphibia: a comparative study with soybean agglutinin. J. Hirnforsch. 33, 627–35. Available at: http://www.ncbi.nlm.nih.gov/pubmed/1494040.

Freitag, J., Krieger, J., Strotmann, J., and Breer, H. (1995). Two classes of olfactory receptors in Xenopus laevis. Neuron 15, 1383–1392. doi:10.1016/0896-6273(95)90016-0.

Freitag, J., Ludwig, G., Andreini, I., Roessler, P., Breer, H., Rössler, P., et al. (1998). Olfactory receptors in aquatic and terrestrial vertebrates. J. Comp. Physiol. A 183, 635–650. doi:10.1007/s003590050287.

Friedrich, J., Zhou, P., and Paninski, L. (2017). Fast online deconvolution of calcium imaging data. PLoS Comput. Biol. 13, e1005423. doi:10.1371/journal.pcbi.1005423.

Gaudin, A., and Gascuel, J. (2005). 3D atlas describing the ontogenic evolution of the primary olfactory projections in the olfactory bulb of Xenopus laevis. J. Comp. Neurol. 489, 403–424. doi:10.1002/cne.20655.

Giovannucci, A., Friedrich, J., Gunn, P., Kalfon, J., Brown, B. L., Koay, S. A., et al. (2019). CaImAn an open source tool for scalable calcium imaging data analysis. Elife 8, e38173. doi:10.7554/eLife.38173.

Gliem, S., Syed, A. S., Sansone, A., Kludt, E., Tantalaki, E., Hassenklöver, T., et al. (2013). Bimodal processing of olfactory information in an amphibian nose: Odor responses segregate into a medial and a lateral stream. Cell. Mol. Life Sci. 70, 1965–1984. doi:10.1007/s00018-012-1226-8.

Hansen, A., Reiss, J. O., Gentry, C. L., and Burd, G. D. (1998). Ultrastructure of the olfactory organ in the clawed frog, Xenopus laevis , during larval development and metamorphosis. J. Comp. Neurol. 288, 273–288. doi:10.1002/(SICI)1096-9861(19980824)398:2<273::AID-CNE8>3.0.CO;2-Y.

Hassenklöver, T., and Manzini, I. (2014). The olfactory system as a model to study axonal growth patterns and orphology in vivo. J. Vis. Exp., e52143. doi:10.3791/52143.

Heerema, J. L., Bogart, S. J., Helbing, C. C., and Pyle, G. G. (2020). Olfactory epithelium ontogenesis and function in postembryonic North American Bullfrog (Rana (Lithobates) catesbeiana) tadpoles. Can. J. Zool. 98, 367–375. doi:10.1139/cjz-2019-0213.

Helling, H. (1938). Das Geruchsorgan der Anuren, vergleichend-morphologisch betrachtet. Z. Anat. Entwicklungsgesch. 108, 587–643. doi:10.1007/BF02118847.

Herrick, C. J. (1924). The amphibian forebrain. II. The olfactory bulb of Amblystoma. J. Comp. Neurol. 37, 55–69. doi:10.1002/cne.900530103.

Higgs, D. M., and Burd, G. D. (2001). Neuronal turnover in the Xenopus laevis olfactory epithelium during metamorphosis. J. Comp. Neurol. 433, 124–130. doi:10.1002/cne.1130.

Hoffman, H. H. (1963). The olfactory bulb, accessory bulb, and hemisphere of some anurans. J. Comp. Neurol. 120, 317–368. doi:10.1002/cne.901200208.

Hofmann, M. H., and Meyer, D. L. (1991). Functional subdivisions of the olfactory system correlate with lectin-binding properties in Xenopus. Brain Res. 564, 344–347. doi:10.1016/0006-8993(91)91475-G.

Hourdry, J., L’Hermite, A., and Ferrand, R. (1996). Changes in the digestive tract and feeding behavior of anuran amphibians during metamorphosis. Physiol. Zool. 69, 219–251. doi:10.1086/physzool.69.2.30164181.

Huang, L. K., and Wang, M. J. J. (1995). Image thresholding by minimizing the measures of fuzziness. Pattern Recognit. 28, 41–51. doi:10.1016/0031-3203(94)E0043-K.

Huber, C. G., and Crosby, E. C. (1929). The nuclei and fiber paths of the avian diencephalon, with consideration of telencephalic and certain mesencephalic centers and connections. J. Comp. Neurol. 48, 1–225. doi:10.1002/cne.900480102.

Ignell, R., Anton, S., and Hansson, B. S. (2001). The antennal lobe of orthoptera - Anatomy and evolution. Brain. Behav. Evol. 57, 1–17. doi:10.1159/000047222.

Imamura, F., Ito, A., and Lafever, B. J. (2020). Subpopulations of projection neurons in the olfactory bulb. Front. Neural Circuits 14, 561822. doi:10.3389/fncir.2020.561822.

Jermakowicz, W. J., Dorsey, D. A., Brown, A. L., Wojciechowski, K., Giscombe, C. L., Graves, B. M., et al. (2004). Development of the nasal chemosensory organs in two terrestrial anurans: The directly developing frog, Eleutherodactylus coqui (Anura: Leptodactylidae), and the metamorphosing toad, Bufo americanus (Anura: Bufonidae). J. Morphol. 261, 225–248. doi:10.1002/jmor.10246.

Jiang, T., and Holley, A. (1992a). Morphological variations among output neurons of the olfactory bulb in the frog (Rana ridibunda). J. Comp. Neurol. 320, 86–96. doi:10.1002/cne.903200106.

Jiang, T., and Holley, A. (1992b). Some properties of receptive fields of olfactory mitral/tufted cells in the frog. J. Neurophysiol. 68, 726–733. doi:10.1152/jn.1992.68.3.726.

Jungblut, L. D., Pozzi, A. G., and Paz, D. A. (2011). Larval development and metamorphosis of the olfactory and vomeronasal organs in the toad Rhinella (Bufo) arenarum (Hensel, 1867). Acta Zool. 92, 305–315. doi:10.1111/j.1463-6395.2010.00461.x.

Jungblut, L. D., Pozzi, A. G., and Paz, D. A. (2012). A putative functional vomeronasal system in anuran tadpoles. J. Anat. 221, 364–372. doi:10.1111/j.1469-7580.2012.01543.x.

Jungblut, L. D., Reiss, J. O., Paz, D. A., and Pozzi, A. G. (2017). Quantitative comparative analysis of the nasal chemosensory organs of anurans during larval development and metamorphosis highlights the relative importance of chemosensory subsystems in the group. J. Morphol., 1–12. doi:10.1002/jmor.20705.

Jungblut, L. D., Reiss, J. O., and Pozzi, A. G. (2021). Olfactory subsystems in the peripheral olfactory organ of anuran amphibians. Cell Tissue Res. 383, 289–299. doi:10.1007/s00441-020-03330-6.

Kapur, J. N., Sahoo, P. K., and Wong, A. K. C. (1985). A new method for gray-level picture thresholding using the entropy of the histogram. Comput. Vision, Graph. Image Process. 29, 273–285. doi:10.1016/0734-189X(85)90125-2.

Key, B., and Giorgi, P. P. (1986). Selective binding of soybean agglutinin to the olfactory system of Xenopus. Neuroscience 18, 507–15. doi:10.1016/0306-4522(86)90171-5.

Laberge, F. (2008). Cytoarchitecture of the accessory olfactory bulb in the salamander Plethodon shermani. Brain Res. 1219, 32–45. doi:10.1016/j.brainres.2008.04.042.

Macrides, F., and Schneider, S. P. (1982). Laminar organization of mitral and tufted cells in the main olfactory bulb of the adult hamster. J. Comp. Neurol. 208, 419–430. doi:10.1002/cne.902080410.

Macrides, F., Schoenfeld, T. A., Marchand, J. E., and Clancy, A. N. (1985). Evidence for morphologically, neurochemically and functionally heterogeneous classes of mitral and tufted cells in the olfactory bulb. Chem. Senses 10, 175–202. doi:10.1093/chemse/10.2.175.

Manzini, I., Rössler, W., and Schild, D. (2002). cAMP-independent responses of olfactory neurons in Xenopus laevis tadpoles and their projection onto olfactory bulb neurons. J. Physiol. 545, 475–84. doi:10.1113/jphysiol.2002.031914.

Manzini, I., and Schild, D. (2003). cAMP-independent olfactory transduction of amino acids in Xenopus laevis tadpoles. J. Physiol. 551, 115–23. doi:10.1113/jphysiol.2003.043059.

Manzini, I., and Schild, D. (2010). “Olfactory coding in larvae of the African Clawed frog Xenopus laevis,” in The Neurobiology of Olfaction, ed. A. Menini (Boca Raton, FL: CRC Press/Taylor & Francis), 114–126. Available at: http://www.ncbi.nlm.nih.gov/books/NBK55981/.

Measey, J. (2016). Overland movement in African clawed frogs (*Xenopus laevis*): a systematic review. PeerJ 4, e2474. doi:10.7717/peerj.2474.

Mezler, M., Fleischer, J., and Breer, H. (2001). Characteristic features and ligand specificity of two olfactory receptor classes from Xenopus laevis. J. Exp. Biol. 204, 2987–2997. doi:10.1242/jeb.204.17.2987.

Mezler, M., Konzelmann, S., Freitag, J., Rössler, P., and Breer, H. (1999). Expression of olfactory receptors during development in Xenopus laevis. J. Exp. Biol. 202, 365–376. doi:10.1242/jeb.202.4.365.

Mombaerts, P. (2006). Axonal wiring in the mouse olfactory system. Annu. Rev. Cell Dev. Biol. 22, 713–737. doi:10.1146/annurev.cellbio.21.012804.093915.

Mori, K., Kishi, K., and Ojima, H. (1983). Distribution of dendrites of mitral, displaced mitral, tufted, and granule cells in the rabbit olfactory bulb. J. Comp. Neurol. 219, 339–355. doi:10.1002/cne.902190308.

Nagayama, S., Homma, R., and Imamura, F. (2014). Neuronal organization of olfactory bulb circuits. Front. Neural Circuits 8, 98. doi:10.3389/fncir.2014.00098.

Nezlin, L. P., Heermann, S., Schild, D., and Rössler, W. (2003). Organization of glomeruli in the main olfactory bulb of Xenopus laevis tadpoles. J. Comp. Neurol. 464, 257–268. doi:10.1002/cne.10709.

Nieuwenhuys, R. (1966). Comparative anatomy of olfactory centres and tracts. Prog. Brain Res. 23, 1–64. doi:10.1016/S0079-6123(08)60662-7.

Nieuwkoop, D. P., and Faber, J. (1994). Normal table of Xenopus laevis (Daudin). Garland, New York.

Nowack, C., and Vences, M. (2016). Ontogenetic development of the derived olfactory system of the mantellid frog Mantidactylus betsileanus. Anat. Rec. 299, 943–950. doi:10.1002/ar.23351.

Offner, T., Daume, D., Weiss, L., Hassenklöver, T., and Manzini, I. (2020). Whole-brain calcium imaging in larval Xenopus. Cold Spring Harb. Protoc. 2020, pdb.prot106815. doi:10.1101/pdb.prot106815.

Orona, E., Rainer, E. C., and Scott, J. W. (1984). Dendritic and axonal organization of mitral and tufted cells in the rat olfactory bulb. J. Comp. Neurol. 226, 346–56. doi:10.1002/cne.902260305.

Pedregosa, F., Varoquaux, G., Gramfort, A., Michel, V., Thirion, B., Grisel, O., et al. (2011). Scikit-learn: Machine learning in python. J. Mach. Learn. Res. 12, 2825–2830. Available at: https://scikit-learn.org/stable/.

Peng, H., Ruan, Z., Long, F., Simpson, J. H., and Myers, E. W. (2010). V3D enables real-time 3D visualization and quantitative analysis of large-scale biological image data sets. Nat. Biotechnol. 28, 348–353. doi:10.1038/nbt.1612.

Pnevmatikakis, E. A., Gao, Y., Soudry, D., Pfau, D., Lacefield, C., Poskanzer, K., et al. (2014). A structured matrix factorization framework for large scale calcium imaging data analysis. arXiv Prepr. arXiv1409.2903.

Pnevmatikakis, E. A., and Giovannucci, A. (2017). NoRMCorre: An online algorithm for piecewise rigid motion correction of calcium imaging data. J. Neurosci. Methods 291, 83–94. doi:10.1016/j.jneumeth.2017.07.031.

Pnevmatikakis, E. A., Soudry, D., Gao, Y., Machado, T. A., Merel, J., Pfau, D., et al. (2016). Simultaneous denoising, deconvolution, and demixing of calcium imaging data. Neuron 89, 285–299. doi:10.1016/j.neuron.2015.11.037.

Preibisch, S., Saalfeld, S., and Tomancak, P. (2009). Globally optimal stitching of tiled 3D microscopic image acquisitions. Bioinformatics 25, 1463–1465. doi:10.1093/bioinformatics/btp184.

Quinzio, S. I., and Reiss, J. O. (2018). The ontogeny of the olfactory system in ceratophryid frogs (Anura, Ceratophryidae). J. Morphol. 279, 37–49. doi:10.1002/jmor.20751.

Reiss, J. O., and Burd, G. D. (1997a). Cellular and molecular interactions in the development of the Xenopus olfactory system. Semin. Cell Dev. Biol. 8, 171–179. doi:10.1006/scdb.1996.0138.

Reiss, J. O., and Burd, G. D. (1997b). Metamorphic remodeling of the primary olfactory projection in Xenopus: Developmental independence of projections from olfactory neuron subclasses. J. Neurobiol. 32, 213–222. doi:10.1002/(SICI)1097-4695(199702)32:2<213::AID-NEU6>3.0.CO;2-B.

Reiss, J. O., and Eisthen, H. L. (2008). “Comparative anatomy and physiology of chemical senses in amphibians,” in Sensory evolution on the threshold: Adaptations in secondarily aquatic vertebrates, eds. J. Thewissen and S. Nummela (Oakland, CA: University of California Press), 43–63. doi:10.1525/california/9780520252783.003.0007.

Sansone, A., Hassenklöver, T., Syed, A. S., Korsching, S. I., and Manzini, I. (2014). Phospholipase C and diacylglycerol mediate olfactory responses to amino acids in the main olfactory epithelium of an amphibian. PLoS One 9, e87721. doi:10.1371/journal.pone.0087721.

Savage, R. M. (1965). External stimulus for the natural spawning of Xenopus laevis. Nature 205, 618–619. doi:10.1038/205618a0.

Scalia, F., Gallousis, G., and Roca, S. (1991). A note on the organization of the amphibian olfactory bulb. J. Comp. Neurol. 305, 435–442. doi:10.1002/cne.903050307.

Schindelin, J., Arganda-Carreras, I., Frise, E., Kaynig, V., Longair, M., Pietzsch, T., et al. (2012). Fiji: an open-source platform for biological-image analysis. Nat. Methods 9, 676–682. doi:10.1038/nmeth.2019.

Scott, D. W. (1992). Multivariate Density Estimation: Theory, Practice, and Visualization. , ed. D. W. Scott Hoboken, New Jersey: John Wiley & Sons, Ltd doi:10.1002/9780470316849.

Sorensen, P., and Caprio, J. (1998). “Chemoreception,” in The physiology of fishes, ed. D. Evans (Boca Raton, FL: CRC), 375–405.

Syed, A. S., Sansone, A., Hassenklöver, T., Manzini, I., Korsching, S. I., Sciences, M. L., et al. (2017). Coordinated shift of olfactory amino acid responses and V2R expression to an amphibian water nose during metamorphosis. Cell. Mol. Life Sci. 74, 1711–1719. doi:10.1007/s00018-016-2437-1.

Syed, A. S., Sansone, A., Nadler, W., Manzini, I., and Korsching, S. I. (2013). Ancestral amphibian v2rs are expressed in the main olfactory epithelium. Proc. Natl. Acad. Sci. 110, 7714–7719. doi:10.1073/pnas.1302088110.

Virtanen, P., Gommers, R., Oliphant, T. E., Haberland, M., Reddy, T., Cournapeau, D., et al. (2020). SciPy 1.0: fundamental algorithms for scientific computing in Python. Nat. Methods 17, 261–272. doi:10.1038/s41592-019-0686-2.

Wakisaka, N., Miyasaka, N., Koide, T., Masuda, M., Hiraki-Kajiyama, T., and Yoshihara, Y. (2017). An adenosine receptor for olfaction in fish. Curr. Biol. 27, 1–11. doi:10.1016/j.cub.2017.04.014.

Weiss, L., Jungblut, L. D., Pozzi, A. G., O’Connell, L. A., Hassenklöver, T., and Manzini, I. (2020a). Conservation of glomerular organization in the main olfactory bulb of anuran larvae. Front. Neuroanat. 14, 44. doi:10.3389/fnana.2020.00044.

Weiss, L., Jungblut, L. D., Pozzi, A. G., Zielinski, B. S., O’Connell, L. A., Hassenklöver, T., et al. (2020b). Multi-glomerular projection of single olfactory receptor neurons is conserved among amphibians. J. Comp. Neurol. 528, 2239–2253. doi:10.1002/cne.24887.

Weiss, L., Manzini, I., and Hassenklöver, T. (2021). Olfaction across the water–air interface in anuran amphibians. Cell Tissue Res. 383, 301–325. doi:10.1007/s00441-020-03377-5.

Weiss, L., Offner, T., Hassenklöver, T., and Manzini, I. (2018). “Dye electroporation and imaging of calcium signaling in Xenopus nervous system,” in Methods in Molecular Biology, ed. K. Vleminckx (New York, NY: Humana), 217–231. doi:10.1007/978-1-4939-8784-9_15.

Wells, K. D. (2007). The ecology and behavior of amphibians. Chicago, IL: University of Chicago Press.

Yabuki, Y., Koide, T., Miyasaka, N., Wakisaka, N., Masuda, M., Ohkura, M., et al. (2016). Olfactory receptor for prostaglandin F2α mediates male fish courtship behavior. Nat. Neurosci. 19, 897–904. doi:10.1038/nn.4314.

Yokosuka, M., Hagiwara, A., Saito, T. R., Aoyama, M., Ichikawa, M., and Sugita, S. (2009a). Morphological and histochemical study of the nasal cavity and fused olfactory bulb of the brown-eared bulbul, Hysipetes amaurotis. Zoolog. Sci. 26, 713–721. doi:10.2108/zsj.26.713.

Yokosuka, M., Hagiwara, A., Saito, T. R., Tsukahara, N., Aoyama, M., Wakabayashi, Y., et al. (2009b). Histological properties of the nasal cavity and olfactory bulb of the Japanese jungle crow corvus macrorhynchos. Chem. Senses 34, 581–593. doi:10.1093/chemse/bjp040.

Yopak, K. E., Lisney, T. J., and Collin, S. P. (2015). Not all sharks are “swimming noses”: variation in olfactory bulb size in cartilaginous fishes. Brain Struct. Funct. 220, 1127–1143. doi:10.1007/s00429-014-0705-0.

